# Selection of HIV-1 for Resistance to Fifth Generation Protease Inhibitors Reveals Two Independent Pathways to High-Level Resistance

**DOI:** 10.1101/837120

**Authors:** Ean Spielvogel, Sook-Kyung Lee, Shuntai Zhou, Gordon J. Lockbaum, Mina Henes, Amy Sondegroth, Klajdi Kosovrasti, Ellen A. Nalivaika, Akbar Ali, Nese Kurt Yilmaz, Celia A. Schiffer, Ronald Swanstrom

## Abstract

Darunavir (DRV) is exceptional among potent HIV-1 protease inhibitors (PIs) in the high drug concentrations that are achived *in vivo*. Little is known about the *de novo* resistance pathway for DRV. We have selected for resistance to high drug concentrations against ten PIs and their structural precursor DRV. Mutations accumulated through two pathways (anchored by protease mutations I50V or I84V). Small chemical changes in the inhibitor P1’ side chain led to preferential use of one pathway over the other. Changes in the inhbitor P2’ side chain determined differences in potency that were retained in the resistant viruses. Viral variants from the two pathways showed differential selection of compensatory mutations in Gag cleavage sites. These results reveal the high level of selective pressure that is attainable with fourth generation PIs, and how features of the inhibitor affect both the resistance pathway and the residual potency in the face of resistance.

## Introduction

Highly active antiretroviral therapy against HIV-1 with combinations of drugs effectively block viral replication and preclude the evolution of drug resistance. Three major factors interplay to define the emergence of resistance *in vivo*: i) the active drug concentration relative to its inhibitory activity; ii) the level of resistance conferred by one or more mutations; and iii) the fitness cost of the resistance mutations. For protease inhibitors (PIs), the number of mutations selected increases with increasing drug concentration (Watkins et al., 2003). Thus the concentration of this drug class *in vivo* is an important parameter in determining the genetic barrier to resistance.

The early identification of the retroviral protease as a member of the aspartyl proteinase family and the determination of a number of cleavage site sequences led to the development of first generation PIs (PIs) that validated the HIV-1 protease as a drug target (Katoh et al., 1987; Richards et al., 1989; Seelmeier et al., 1988). A second generation of PIs was quickly developed for use in humans, becoming the third drug in a three-drug regimen that achieved sustained suppression of viral load with no evolution of resistance (Gulick et al., 1997). The third generation of PIs had improved properties with regard to side effects and efficacy. These properties have been further enhanced with a fourth-generation PI, darunavir (DRV), which achieves a drug level in plasma (>1 µM) that is 1,000 fold greater than its inhibitory activity in cell culture (Ali et al., 2010; Nalam et al., 2013; Kurt Yilmaz et al., 2009). The potential efficacy of a fourth-generation PI such as DRV has led to attempts to use this drug in monotherapy (Antinori et al., 2015; Arribas et al., 2012; Paton et al., 2015; Valantin et al., 2012). Although the initial trials of monotherapy showed a modest increase in the loss of viral suppression relative to standard ART, in the cases of virologic failure there was no significant resistance to DRV detected in the rebound virus (Antinori et al., 2015; Arribas et al., 2012; Paton et al., 2015; Valantin et al., 2012). Thus the observed virologic failure is most easily attributed to issues of low drug exposure in specific participants. While therapy has not yet moved to single agents, potent two drug therapies are being explored. In this regard, several studies have shown that a PI can be part of a successful two drug therapy regimen (Casado et al., 2020; Di Cristo et al., 2020; Hawkins et al., 2019; Huang et al., 2019; Vizcarra et al., 2020).

Drug resistance to DRV is incompletely understood. The appearance of DRV-resistant viruses during therapy failure has occurred in people who had previously experienced virologic failure with other PIs (de Meyer et al., 2008; Descamps et al., 2009; Pellegrin et al., 2008; Tremblay, 2008). In these cases the pathway to DRV resistance was likely influenced by the pre-existing resistance to the previous PI. DRV resistance has been selected in cell culture (Aoki et al., 2018; de Meyer et al., 2008; Koh et al., 2010). High level resistance was selected starting with either a mixture of highly resistant variants that were allowed to recombine, or the selection was started with a single PI-resistant strain; in both cases an I84V resistance pathway in the protease was observed. However, there are no reports of selection of high level resistance starting with an unmutated “wild type” virus. Thus there is little information on what a *do novo* pathway for resistance to DRV would be.

An important concept in HIV-1 PI design is to avoid chemical moieties that extend outside of the substrate envelope, the shared volume in the protease active site that is occupied by natural protease substrates (King et al., 2004; Prabu-Jeyabalan et al., 2002). We have previously designed a series of highly potent PIs based on DRV, UMASS-1 through −10, that still fit within the substrate envelope (Nalam et al., 2013). The designed inhibitors have modified chemical moieties that relative to DRV further fill the substrate envelope at the equivalent of the P1’ and P2’ positions. All bind tightly to the wild type HIV-1 protease with a Ki of less than 5 pM. These inhibitors retained robust binding to many multi-drug resistant protease variants and viral strains.

In an effort to define the *de novo* pathway to resistance for DRV and determine the effects of these chemical changes in the inhibitors, we examined the evolutionary path used to attain high level resistance to DRV and the UMASS-1 through −10 panel of PIs. Under continuous and escalating selective pressure, representing between 50-95 passages, the virus evolved to accumulate multiple mutations. In most cases, selection was carried out until the the final concentration approximated that achieved by DRV *in vivo*. Relatively minor modifications in inhibitor structure favored selection of one of two pathways to resistance, anchored by protease mutation I50V or I84V. In addition, one side chain of the inhibitor increased the residual potency against highly resistant strains. These results reveal the extremely high genetic barrier to resistance to fourth-generation PIs at inhibitor concentrations that can be achieved *in vivo*, and the complex evolutionary pathways required to achieve resistance. In addition, these results reveal features of inhibitor design that both affect the selected pathway and determine residual potency against resistant variants.

## Results

### Panel of highly potent and analogous HIV-1 PIs

HIV-1 PIs were designed by modifications to DRV to increase favorable interactions with the protease within the substrate envelope thereby increasing potency while minimizing evolution of resistance (Nalam et al., 2013). A panel of ten DRV analogues were chosen with enzymatic inhibition constants (K_i_) in the single or double-digit picomolar range to wild-type NL4-3 protease and the I84V and I50V/A71V drug resistant variants, respectively (Table 1) (Lockbaum et al., 2019; Mittal et al., 2013). These PIs contained modified P1’ positions with (S)-2-methylbutyl or 2-ethyl-n-butyl moieties (R1-1 and R1-2, respectively) in combination with five diverse P2′ phenyl-sulfonamides (R2-1 to R2-5), with the inhibitors named UMASS-1 through −10 (Table 1). These inhibitors and DRV were also tested in a cell culture-based viral inhibition assay. The EC_50_ values (the effective concentration needed at the time of virus production to reduce infectivity by 50%) for DRV and the UMASS analogues ranged from 2.4 to 9.1 nM, significantly more potent than the second and third generation PIs tested (Fig. S1).

**Table 1.** Structures, Ki Constants, and EC50 Values of the UMass Series of HIV-1 PIs

| Inhibitor | Structure | Ki (pM) |  |  | EC50 (nM) |
| --- | --- | --- | --- | --- | --- |
|  |  | WT | I84V | I50V/A71V | WT |
| DRV |  | < 5.0<sup>ψ</sup> | 25.6 ± 5.6<sup>a</sup> | 74.5 ± 5.6<sup>a</sup> | 7.7 |
| UMASS-1 |  | < 5.0<sup>ψ</sup> | 26.1 ± 3.7<sup>a</sup> | 110.3 ± 8.8<sup>a</sup> | 5.9 |
| UMASS-2 |  | < 5.0 | < 5.0 | 15.0 ± 2.7 | 2.4 |
| UMASS-3 |  | < 5.0 | 9.9 ± 2.7 | 79.9 ± 5.9 | 9.1 |
| UMASS-4 |  | < 5.0 | 10.5 ± 1.8 | 32.9 ± 3.0 | 3.2 |
| UMASS-5 |  | < 5.0 | 7.0 ± 1.7 | 7.8 ± 0.9 | 4.0 |
| UMASS-6 |  | < 5.0<sup>ψ</sup> | 12.8 ± 3.1<sup>a</sup> | 100.0 ± 9.9<sup>a</sup> | 5.2 |
| UMASS-7 |  | < 5.0 | 12.1 ± 4.5 | 18.2 ± 3.0 | 3.1 |
| UMASS-8 |  | < 5.0 | < 5.0 | 55.4 ± 4.0 | 4.2 |
| UMASS-9 |  | < 5.0 | 7.6 ± 1.6 | 42.3 ± 2.6 | 6.4 |
| UMASS-10 |  | < 5.0 | 14.3 ± 9.3 | 5.8 ± 1.1 | 4.1 |

<sup>a</sup>Previously reported in Lockbaum et al (ACS Infect Dis. 2019 Feb 8; 5(2): 316–325.)

There are several patterns worth noting in the K_i_ and EC_50_ values. First, all inhibitors had a K_i_ of less than 5 pM (limit of detection) against the wild type enzyme, precluding a comparison of potency among them without using partially resistant mutant proteases (I84V and I50V/A71V). Second, the single P1’/R1 position change to the larger 2-ethyl-n-butyl moiety (R1-2, UMASS-6) increased potency against the I84V mutant protease relative to DRV while the intermediate sized R1-1 moiety did not (UMASS-1). Third, four of the inhibitors with the R2 moieties R2-2 or R2-5 were significantly more potent (UMASS-2, −5, −7, −10) than with the other R2 moieties. This trend was most pronounced for activity against the I50V/A71V mutant enzyme in both the R1-1 and R1-2 backgrounds, and for the I84V mutant enzyme in the R1-1 background (Fig. S2). The significant potency of the R2-2 and R2-5 containing inhibitors against the I50V/A71V mutant enzyme while maintaining good potency against the I84V enzyme indicates that both the R1 and R2 constituents contribute to potency, and to enhancing potency over DRV. This potency was also seen in inhibition of viral infectivity as the EC_50_ values of the R2-2 and R2-5 containing inhibitors were among the lowest measured (Table 1).

### Selection for high level resistance during passage in cell culture follows two pathways

To evaluate the potential of each inhibitor to select for mutations that would confer high-level resistance and to compare these mutations across the inhibitor series, we grew HIV-1 under conditions of escalating inhibitor concentration in cell culture through a lengthy period of selection (50-95 passages). The selection experiments were performed under two separate starting conditions: first when the starting virus was generated from the NL4-3 clone (this clone closely approximates the clade B consensus for the protease amino acid sequence and will be referred to as “wild type”), then again when the starting virus was a mixture of 26 isogenic viruses each with a single mutation associated with drug resistance in the NL4-3 background. Notably, in the latter case only about one-half of the mutations that appeared in the culture during selections were in the mixture of starting mutations, indicating that even in the selections that were seeded with the pool of single resistance mutations there was sufficient evolutionary capacity to explore mutations at positions that started as the wild type sequence. The initial inhibitor concentration started at low nanomolar concentrations and increased by a factor of 1.5 with each subsequent viral passage (the inhibitor concentration was increased only after the virus spread efficiently through the culture). All of the selections starting with wild type virus reached at least 5 µM of inhibitor concentration (approximating the therapeutic concentration reached by DRV in the blood and in the range of a 1,000 fold increase in the starting EC_50_). For technical reasons, only 5 of the selections starting with the mixture of mutants reached an inhibitor concentration as high as 400 nM and are reported here (Fig. S3).

Resistance mutations selected in the protease coding domain during the escalating selective pressure of increasing PI concentration were examined at various time-points using a next generation sequencing (NGS) protocol that included Primer ID with the MiSeq platform (Zhou et al., 2015). We first examined the sequence of the most abundant variant in each of the two selection schemes present at the highest inhibitor concentration reached (Fig. 1A). For those selections that reached 5 μM, 8 to 14 mutations were present in the most abundant variant. Typically the next two most abundant variants in the population differed by a single amino acid, and on average the three most abundant variants in the 5 μM cultures accounted for 88% of the total viral population (Fig. S4). Evolution of resistance followed two pathways, one based on I50V and one based on I84V. In three cultures, V2wt, V5wt, and V2mut (which reached either 4 or 5 μM final drug concentration; V2wt indicates the virus pool selected with UMASS-2 starting with wild type virus), these two mutations, I84V and I50V, became linked on the same genome (Fig. 1A) showing they are not mutually exclusive.

**Fig. 1.**
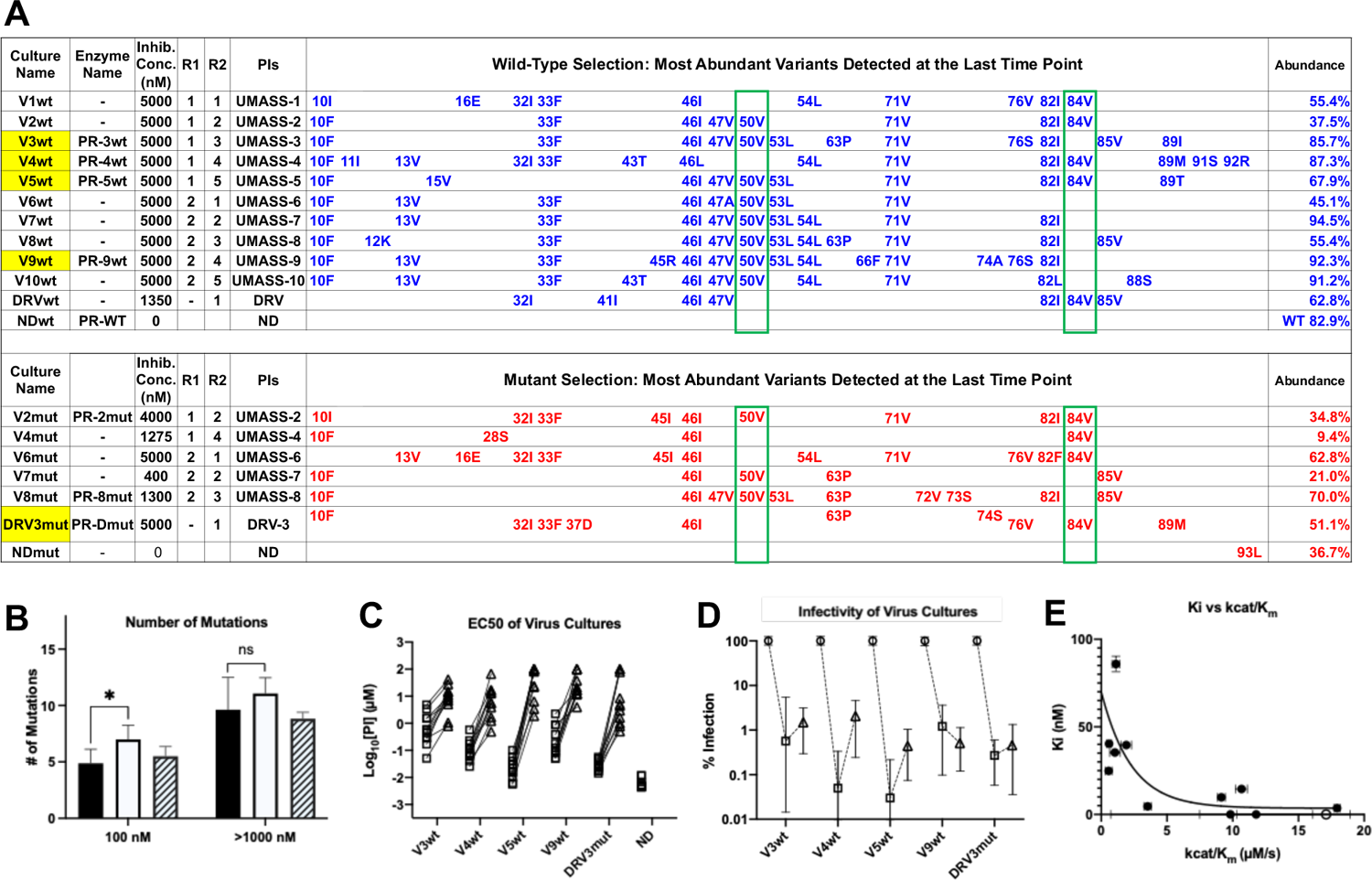
Features of Viruses Selected To High Level Resistance Against DRV and the UMASS-1 Through −10 Inhibitor Series. Infected cultures were put under drug selection starting with either wild type virus or a pool of mutant viruses. **A)** Most abundant variants present at final timepoint. Resistance mutations in the viruses selected starting with the wild type virus are in blue, and the selection starting with the pool of mutant viruses in in red. The culture name indicates a virus pool (V), the UMASS PI number (V1) and whether the selection started with wild type or mutants (V1wt). PR sequences that were used to generate purified recombinant protease are indicated (e.g. PR-3wt). The final PI concentration reached and the R1 and R2 moieties are indicated along with the PI. **B)** Number of mutations present in the 3 most abundant variants of each selection at two different PI concentrations. Cultures containing I84V resistance mutation are in black (n=4), I50V-containing cultures are in white (n=7), and cultures with I50V+I84V linked are shown with hatched box (n=3). Unpaired t-test was used to compare the number of mutations in the 84V vs the 50V cultures. **C)** EC_50_ values for highlighted selections in panel A reaching 100 nM (square) and 5000 nM (triangle). **D)** Relative infectivity values for the same selections shown relative to the wild type virus (circle). Relative infectivity was measured with normalized input amounts of HIV-1 p24 CA protein. **E)** Enzyme inhibition constants (K_i_) of end-point PR variants versus catalytic efficiency (kcat/K_m_). Open circle represents wildtype NL4-3. Trendline is for visualization purposes only. * p <0.05, ** p < 0.01, *** p < 0.001.

In addition to the sequence of the terminally selected virus, we validated that selection occurred over the entire course of the selection protocol. First, as can be seen in Fig. 1B, the number of mutations present at 100 nM drug concentration was approximately half the number present after selection to greater than 1 μM. In comparing the number of mutations present in cultures using the I84V versus the I50V pathway, we noted that on average there were more mutations in the I50V viruses at the intermediate drug concentration compared to the I84V viruses (100 nM, P=0.03), and this trend continued at the higher drug concentration. Next we compared the EC_50_ of five virus cultures representing the I84V or the I50V pathways (highlighted in yellow in Fig. 1), and including one that had both mutations linked, testing the level of resistance to all of the UMASS-1 through −10 inhibitors after selection to 100 nM drug concentration or to 5 μM. As can be seen in Fig. 1C, resistance (measured as an increase in the EC_50_) was apparent after selection to the 100 nM inhibitor concentration, and in each case the EC_50_ increased an average of 100-fold after selection to the final inhibitor concentration of 5 μM.

We were also interested in the fitness cost of these mutations. To measure this effect we compared the infectivity of the virus in 10 culture supernatants (5 selections at each of two levels of selection), normalizing infectivity on a reporter cell line for a given amount of the virion p24 CA protein in the culture supernatant. This was compared to the unmutated NL4-3 parent (given a value of 100%). As can be seen in Fig. 1D, selection to 100 nM inhibitor concentration resulted in a reduction in fitness/relative infectivity by between 100 and 1000-fold. Those cultures with the biggest decrease in fitness rebounded to around 1% relative infectivity after selection to an inhibitor level of 5 μM. This suggests that these high levels of resistance are linked to maintaining a residual level of relative infectivity around 1% (as defined under these culture conditions).

We used another approach to explore the interplay between resistance and fitness; we examined the effect of resistance on catalytic efficiency of the protease by measuring K_i_ and the catalytic efficiency (kcat/Km) for a set of mutant proteases against the entire panel of UMASS inhibitors and DRV. The average K_i_ value of each mutant protease (sequences shown in Table S1) is plotted against the catalytic efficiency (kcat/K_m_) for that enzyme in Fig. 1E. As can be seen, there is a strong relationship between higher K_i_ values (i.e. resistance) and dramatic reductions in catalytic efficiency, consistent with the loss of fitness associated with these viral populations.

### Features of the selection process assessed by next generation sequencing (NGS) of longitudinal samples

Each of the viral cultures started with wild type virus showed an accumulation of protease mutations with increasing selective pressure. NGS analysis revealed very few fixed variants in the cultures started with wild type virus until the inhibitor concentration reached approximately 3-5 nM (approaching the EC_50_ value for the wild type virus, Table 1); in contrast the cultures that were started with the mutant library selected for the outgrowth of a subset of those mutants by 1 nM inhibitor concentration (Fig. 2A). In all cultures, the transition through the EC_50_ concentration provided significant selective pressure in fixing mutations. Multiple unfixed mutations were observed in each culture after the drug concentration exceeded the EC_50_ values above 3 nM, highlighting the high genetic diversity in the culture. Additional mutations became linked on each viral genome at higher drug concentrations.

**Fig. 2.**
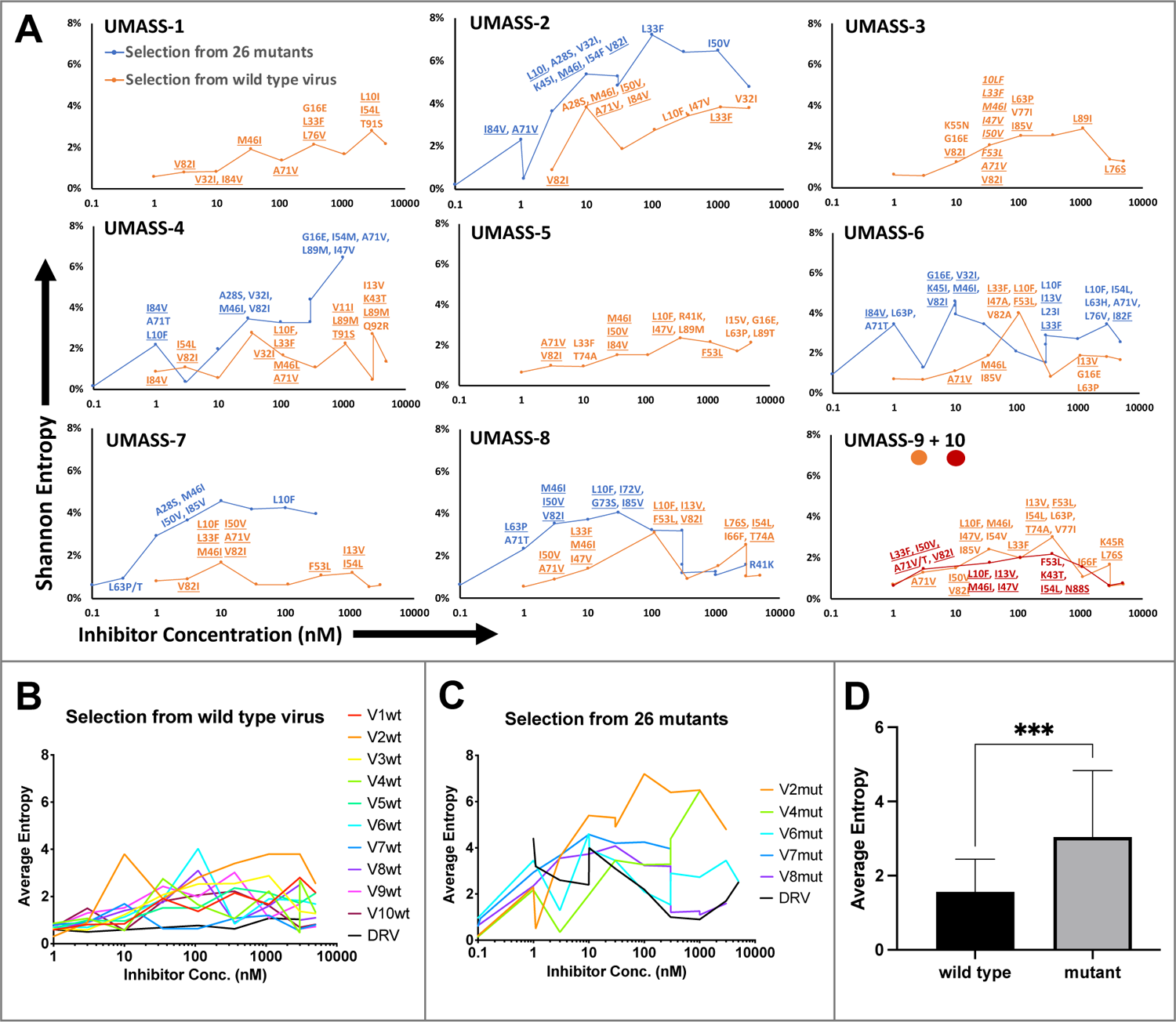
Evaluation of Sequence Diversity In the Viral Population During Increasing Selective Pressure With PI Inhibitors. **A)** Shannon entropy was calculated using sequence diversity in the protease region. Abundant mutations are shown and fixed mutations are underlined. Selections from wild type are in orange (n=6) and from the mutant pool (n=11) in blue. Two selections from wild type virus are shown in orange (UMASS-9) and red (UMASS-10). **B)** Compiled Shannon entropy values from selections starting with wild type virus that reached concentrations of >1000 nM. **C)** Compiled Shannon entropy values from selections starting with the pool of mutant viruses that reached concentrations of at least 400 nM. **D)** Selections starting with the pool of 26 mutants showed higher entropy when averaged over all analyzed drug concentrations (p <0.001 using the unpaired t-test).

Deep sequencing revealed that mutations accumulated in complex patterns. We assessed the sequence complexity of each culture by calculating the Shannon Entropy to allow comparison of changes in the diversity in each culture over time and as a function of increasing selective pressure (Fig. 2A). In the cultures that showed the early appearance of the I84V mutation, this was associated with a peak in entropy, reflecting high genetic diversity, followed by a decrease in entropy when the I84V mutation became fixed (Fig. S5A). The introduction of the I50V mutation was generally not associated with a drop in entropy, rather these populations maintained high genetic diversity even at higher drug concentrations (Fig. S5B). We interpret these patterns as indicative of I84V conferring some level of resistance without a dramatic loss in fitness, allowing a more homogeneous culture (i.e. less entropy). In contrast, I50V may confer a higher level of resistance but at a greater fitness cost thus supporting greater diversity in the culture either as compensatory mutations or as other combinations of mutations with lesser resistance but higher fitness. In this regard, we previously showed I50V significantly reduces the fitness of the virus relative to the fitness loss of a virus with I84V as single mutations (Henderson et al., 2012).

When we examined the sequence diversity as assessed by Shannon Entropy for all selections that reached at least 1 µM in inhibitor concentration, we found that cultures starting with the mixture of resistant viruses averaged a nearly two-fold higher entropy value compared to the cultures where the selection started with just the virus generated from the NL4-3/wild type clone (3.0 vs 1.6, P<0.0001 Mann-Whitney test; Fig. 2B, C, D). This was unexpected, as both sets of selections passed through many genetic bottlenecks. This result is most easily explained if the rates of recombination were fairly high throughout the culture period so that most sequence variants were maintained at least at a low level.

### The chemical nature of R1, but not R2, determines the resistance pathway

We next examined if the inhibitor structure influenced the resistance pathway chosen. We found that the R1 group, i.e. the (S)-2-methylbutyl (R1-1) or the 2-ethyl-n-butyl (R1-2), largely defined the resistance pathway observed; for these inhibitors the R1 group takes the position of P1’ in the protease substrate analog, occupying the S1’ subsite. With the UMASS-1 through −5 series (the smaller R1-1 group), the I84V mutation appeared first in 6 of 7 cultures. In contrast, the UMASS-6 through −10 series with the larger R1-2 group, the I50V mutation appeared first in 8 of 9 cultures (P=0.009, Fisher’s Exact Test; in this analysis we included two cultures that did not reach at least 400 nM inhibitor concentration but did fix an initial set of mutations to increase our sample size, and we did not include two cultures where both I84V and I50V were initially fixed together [data not shown]). We considered the possibility that the starting mixture of mutant viruses in the first selection might skew the pathway selected, especially since the mutant pool included I84V but not I50V. However, in only 1 of the 8 cultures with sufficient data from both selections was there a switch from the I84V pathway to the I50V pathway between the first and second selections (cultures of UMASS-6 with an R1-2 group). Thus, we conclude that the P1’ side chain of the inhibitor is a strong determinant of the resistance pathway selected. It is notable that one inhibitor could select for different pathways in two separate selections (also seen with DRV, see below) even when both major mutations are maintained in the viral population (Fig. S6). This suggests that either pathway can provide some level of resistance to most if not all of these inhibitors, and that the chance addition of the initial compensatory mutations may determine which pathway becomes the major resistant population.

To examine the potential for linked mutations and to infer the order in which mutations accumulated in the protease gene to confer high level resistance, the abundance data from multiple selections that ended in one or the other pathway were pooled and compared. In this analysis, shown in Fig. 3, summary data for the selections resulting in the I84V pathway point up, with I84V reaching 100% penetrance by definition. Similarly, the summary data for those selections that fixed I50V are shown pointing down, with I50V reaching 100% penetrance. Several mutations are uniquely linked or at least strongly favored in each pathway, with I84V being linked to V32I, and I50V being linked to I47V, F53L, and I13V. A number of mutations appear in both pathways, although not with equal frequency: L10F, L33F, M46I, I54L, A71V, and V82I. Finally, other mutations appear less frequently making it difficult to assign them to one of these categories. These results show that while some mutations are largely linked to one pathway, other mutations are often shared between the two pathways. Also, the variation in frequency of appearance of shared mutations in the two pathways suggests different levels of impact on resistance and/or fitness in the I84V vs the I50V background for these mutations (e.g. L10F, M46I and A71V). In particular, the larger impact of I50V on fitness compared to I84V mutations (Henderson et al., 2012) is consistent with the earlier appearance and greater levels of fixation of the shared compensatory mutations L10F, L33F, M46I and A71V in the cultures that followed the I50V pathway. We next considered the possibility that the R2 constituent would provide additional selective pressure in the form of additional resistance mutations; however, similar analyses did not reveal a significant pattern of sequence selection based on the potency associated with the R2 group (data not shown).

**Fig. 3.**
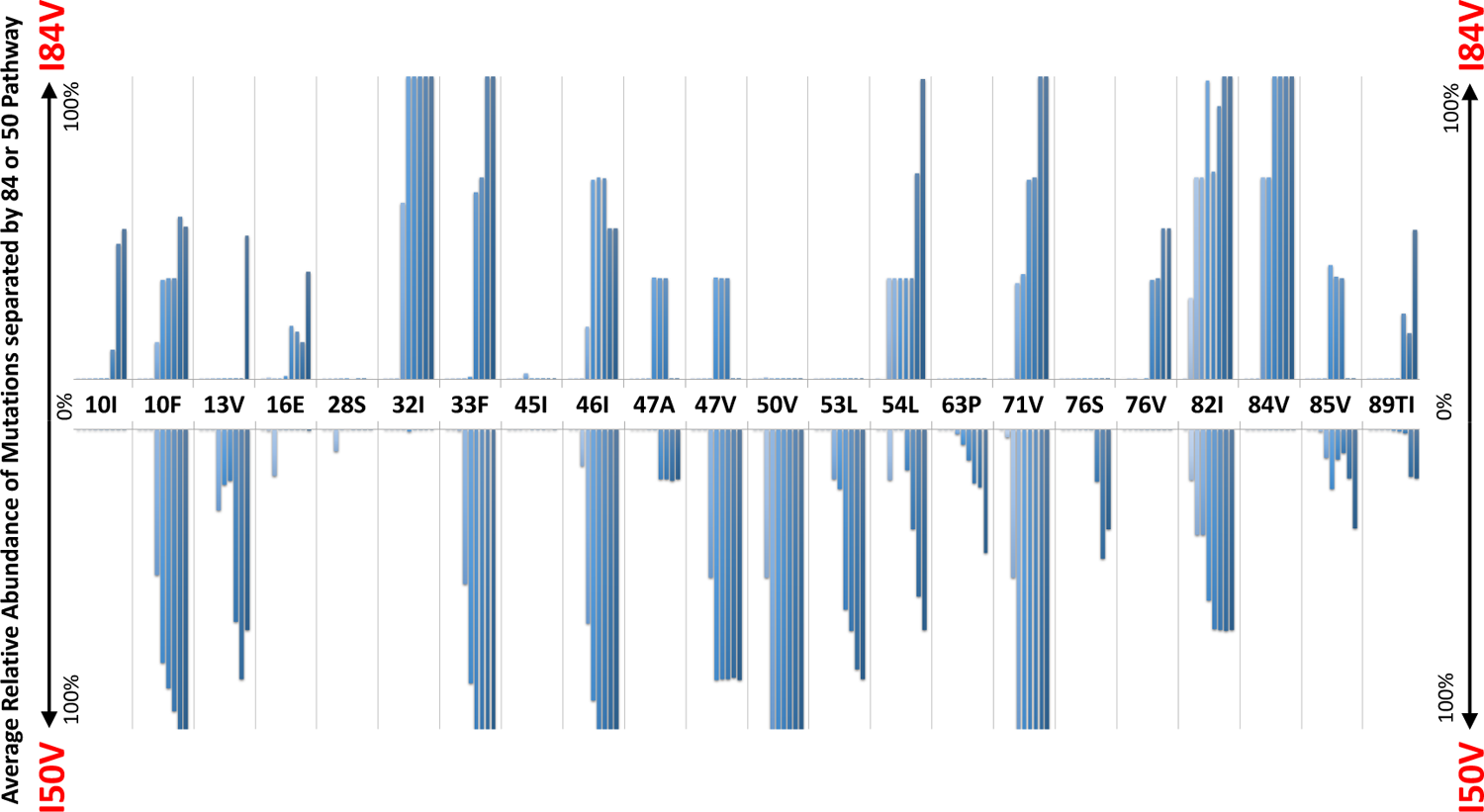
Patterns of Accumulation of Amino Acid Substitutions Associated with Resistance During Selection. Abundance data from selections for each indicated amino acid substitution were pooled and examined sequentially at different levels of drug concentration. Selections that reached the maximum concentration of at least 3000 nM were assessed longitudinally to allow for all timepoints to be averaged equally. Mutations from selections resulting in the I84V pathway (V1wt, V4wt, V6mut, and DRV3mut) point up, with I84V reaching 100% penetrance by definition. Similarly, those selections that fixed I50V only (V3wt, V6-10wt) are shown pointing downward, with I50V reaching 100% penetrance.

### The chemical nature of R1 and R2 determine residual potency among the resistant variants

We considered several variables in examining the nature of the interaction between inhibitors and the resistant proteases: the extent to which the inhibitor structure (either R1 or R2) affected residual potency; and the extent to which the pathway (I84V or I50V) conferred the greatest resistance. As shown in Fig. 4A, the inhibitors with the smaller R1-1 group showed a trend to be less potent than the inhibitors with the larger R1-2 group when tested against all of the resistant proteases. Inhibitors with the R2-2 group showed a trend toward greater residual potency against the mutant enzymes compared to the other inhibitors (Fig. 4B). In looking at the individual pathways, the highly resistant enzymes with the I84V mutation remained more sensitive to the entire group of inhibitors than the enzymes with the I50V mutation (Fig. 4C).

**Fig. 4.**
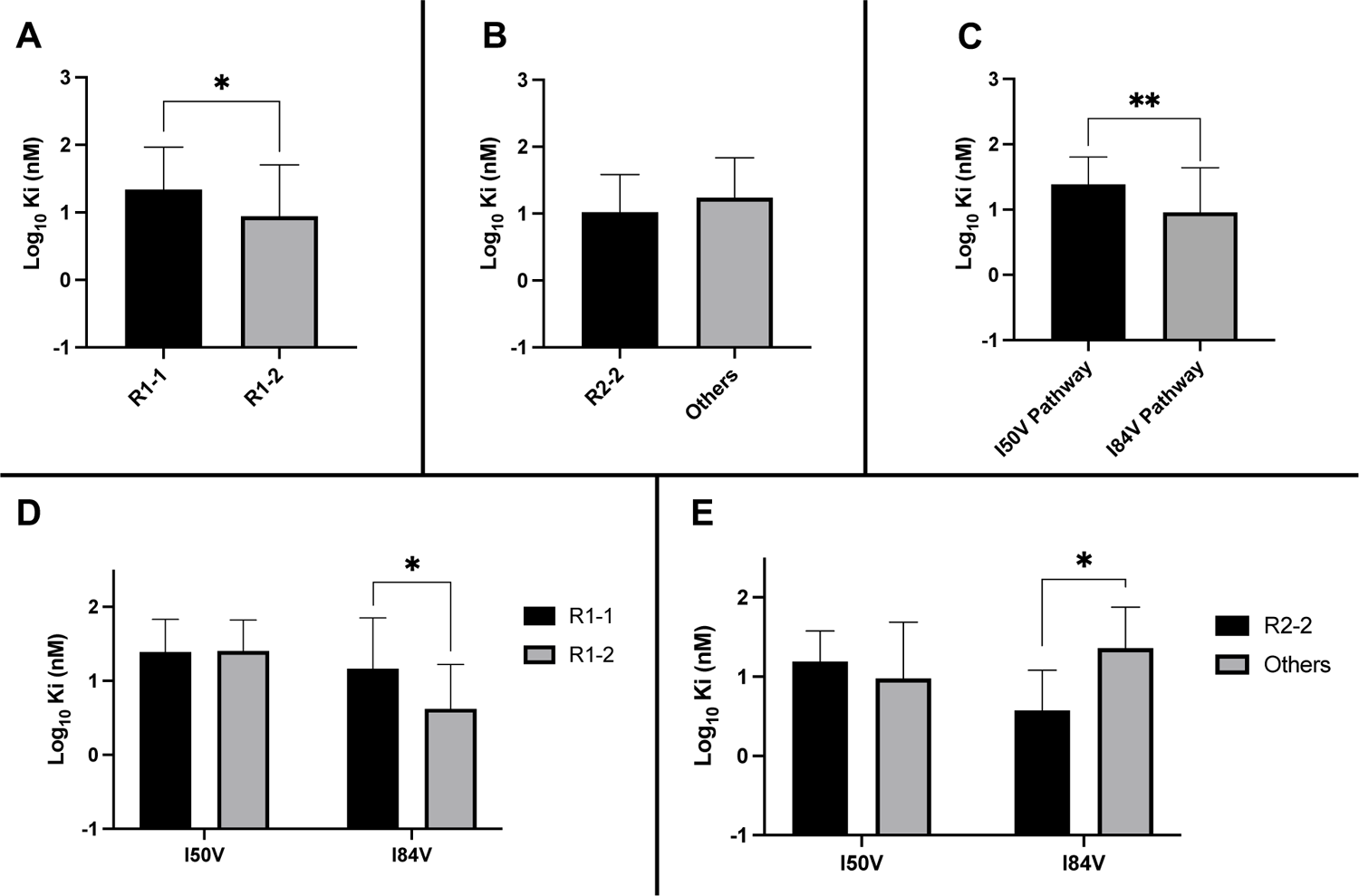
Analysis of Ki Values For Mutant Enzymes. Ki values were determined against all UMASS inhibitors (Table 1). Brackets above the bars represent significant p-values between the two groups using the unpaired t-test. Data were pooled in different ways for the following analyses: **A)** The Ki values against the inhibitors with the larger R1-2 moiety (n=34) were more potent against resistant proteases compared to the inhibitors with the smaller R1-1 moiety (n=35) **B)** Ki values for the inhibitors with the R2-2 group (n=10) showed a trend toward being more potent against the highly mutated proteases compared to the other R2 groups (n=40). **C)** Ki values for enzymes with the I50V mutation (n=33) showed greater resistance to the inhibitors compared to enzymes with the I84V mutation (n=22). **D)** The R1-2 moiety provided increased potency to enzymes with the I84V mutation (n=20) but not the I50V mutation (n=20). **E)** The R2-2 moiety was more potent against the enzymes with I84V mutation (n=10) compared to the enzymes with the I50V mutation (n=10). The unpaired t-test was used to assess differences in K_i_ values.

We next linked the two pathways to the specific structural features of the inhibitors. The resistant proteases with I50V had similar K_i_ values to both the R1-1 and R1-2 inhibitors, while the proteases with I84V were more sensitive (lower K_i_) to the larger R1-2 inhibitors (Fig. 4D). Similarly, the increased potency of the R2-2 inhibitors over the rest of the inhibitors was seen against the enzymes carrying I84V but not those with I50V (Fig. 4E). These results are consistent with the smaller R1-1 inhibitors selecting for the I84V pathway and the larger R1-2 inhibitors selecting for the I50V pathway, and with the R2-2 inhibitors adding additional potency that is retained even after selection for resistance, most apparent with the I84V pathway.

Residual potency dependence of inhibitor structure could also be seen in the EC_50_ values of the resistant virus pools. When the EC_50_ values against the virus pools using the I84V or the I50V pathway were compared for all of the inhibitors, the inhibitors with the larger R1-2 group were more potent, retaining on average lower EC_50_ values compared to the EC_50_ values for the inhibitors with the smaller R1-1 group (Fig. 5A). Inhibitors with the R2-2 group showed a trend toward greater potency compared to the other R2 groups (Fig. 5B), consistent with what was observed in enzymatic assays. When we considered the effects based on the selection pathway we observed that viruses in the I50V pathway had a higher level of resistance to these inhibitors than the viruses in the I84V pathway (Fig. 5C). The higher level of resistance for the viruses using the I50V pathway was due to the fact that these viruses were similarly resistant to the inhibitors with either the R1-1 or the R1-2 groups; in contrast, the viruses using the I84V pathway conferred a greater level of resistance to the inhibitors with the smaller R1-1 group while the inhibitors with the larger R1-2 group retained a higher level of potency (Fig. 5D). Finally, there was a trend for viruses in either pathway to be less resistant to the inhibitors with the R2-2 group (Fig. 5E).

**Fig. 5.**
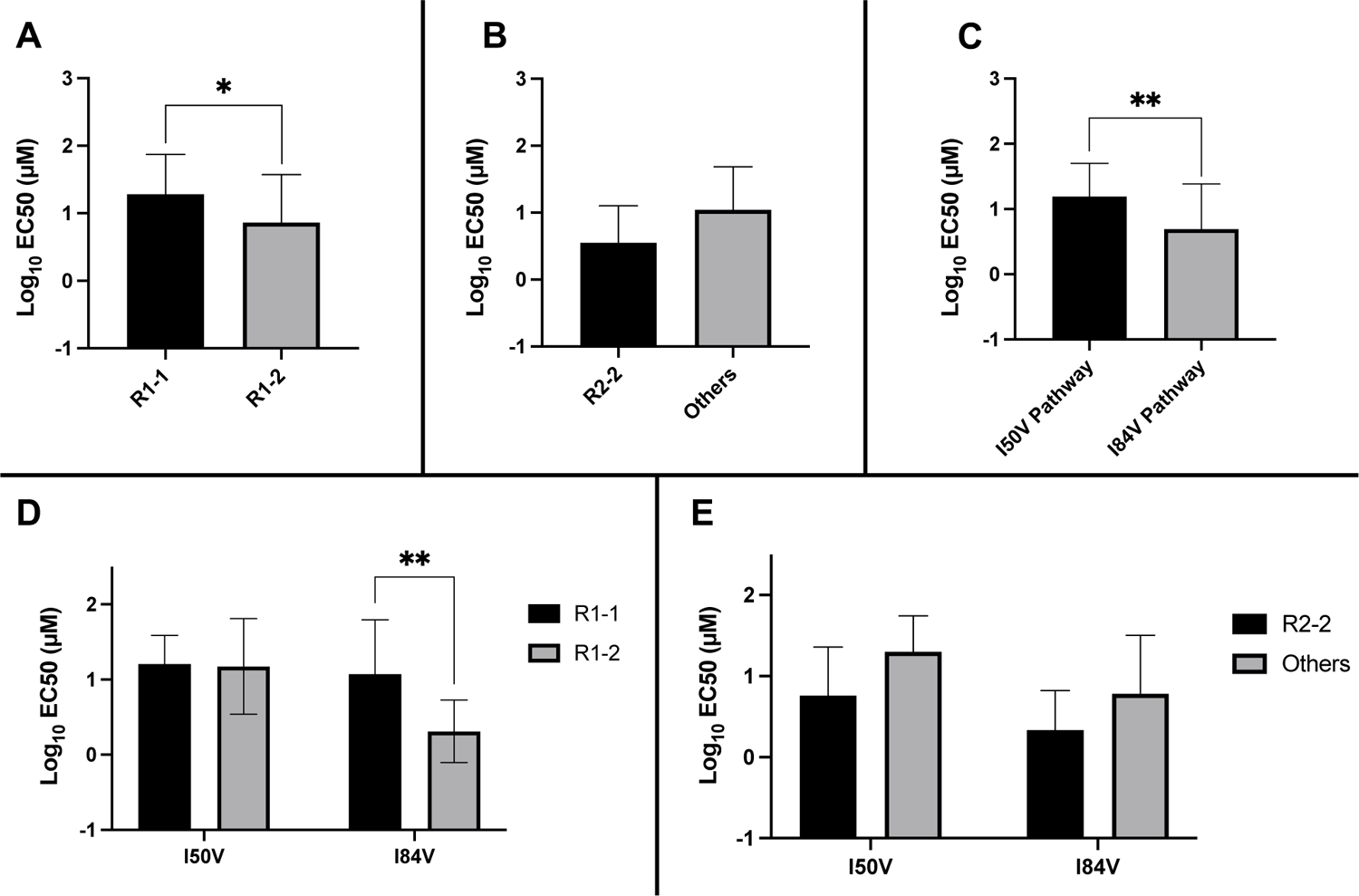
Analysis of EC_50_ Values For Mutant Virus Cultures. EC_50_ values were determined for a subset of the selected virus cultures against a panel of inhibitors (n=50). For **(A-E),** the EC_50_ data were pooled using the same methods as in Fig. 4. The highest level of resistance recorded was 100 µM. The Mann-Whitney rank-sum test was used to assess differences in EC_50_ values.

### DRV favors the I84V pathway

DRV has a butyl group at the R1 position, smaller than the R1-1 methylbutyl in the UMASS-1 through −5 series. We carried out five selections with DRV where the final inhibitor concentration reached greater than 1 μM. One DRV selection was carried out in parallel with each of the two different selections with the UMASS inhibitors (Fig. 1A); both of these selections resulted in the appearance of the I84V mutation as defining the resistance pathway. Three additional selections were done in parallel with DRV and starting with the mixture of the 26 isogenic mutants; all three selections reached the level of 5 μM as the highest drug concentration. Two of these selections used the I84V pathway to resistance, while one of the selections used the I50V pathway to resistance (data not shown). Thus four of five independent selection with DRV favored the resistance pathway I84V, associated with a smaller R1 constituent. These results add further evidence for the size of the R1 moiety strongly influencing the resistance pathway.

## Discussion/Structural Interpretation

DRV is a notable PI both for its tight binding to the HIV-1 protease and for its ability, with boosting doses of ritonavir, to reach micromolar levels of drug concentration in the blood. These features are important as resistance to PIs typically require multiple mutations affecting both resistance and the rescue of enzyme activity/fitness lost with primary resistance mutations (Arribas et al., 2012; Clemente et al., 2004; Henes et al., 2019; Mahalingam et al., 2002; Muzammil et al., 2003; Ragland et al., 2014; Wensing et al., 2010). While we were successful in selecting for resistance to DRV in cell culture, the spectrum of mutations observed largely overlap those seen in previous studies based on testing individual mutations for their effect on DRV sensitivity (de Meyer et al., 2008; King et al., 2004; Rhee et al., 2003). In this work, we have explored chemical modifications to the DRV scaffold. We found that modification of the P1’/R1 chemical moiety to be a larger aliphatic group favored the use of the I50V resistance pathway compared to the smaller DRV moiety or an intermediate-sized moiety, forcing the virus to use the more deleterious I50V mutation compared to the I84V mutation used with the smaller P1’ moieties. Conversely, we identified a subset of the P2’/R2 groups that contributed to residual potency even in the resistant proteases but did not select for additional resistance mutations. Thus chemical modifications at both of these inhibitor positions yielded improvements over DRV.

### P1’/R1 Modification

DRV has a relatively small butyl group at this position that most often selected for an I84V mutation. When we extended this moiety by a single methyl group to (S)-2-methylbutyl we obtained a similar resistance selection profile. However, by extending it with an additional methyl group to (S)-2-methylbutyl or 2-ethyl-n-butyl created an inhibitor that switched the preferred selection to the I50V pathway. In Fig. 6A-C are representations of structures of a subset of the inhibitors with the different P1’/R1 groups with the wild type protease, the I84V mutant protease, and the I50V mutant protease. As we have previously discussed (Lockbaum et al., 2019), in these structures it is apparent that the P1’/R1-1 group is directed at the protease I84 sidechain, consistent with the shortening of this sidechain as a major resistance mechanism. Conversely, the longer P1’/R1-2 group is oriented into the space between I84 and I50, thus drawing in I50V as the pathway to resistance. As noted in the longitudinal analysis of the selection pathway, the I50V resistance pathway presents additional challenges to the virus in the more rapid accumulation of compensatory mutations (Fig. 3). Under the circumstances of rapidly declining viral load during the initiation of therapy this need for additional mutations would represent an enhanced genetic barrier.

**Fig. 6.**
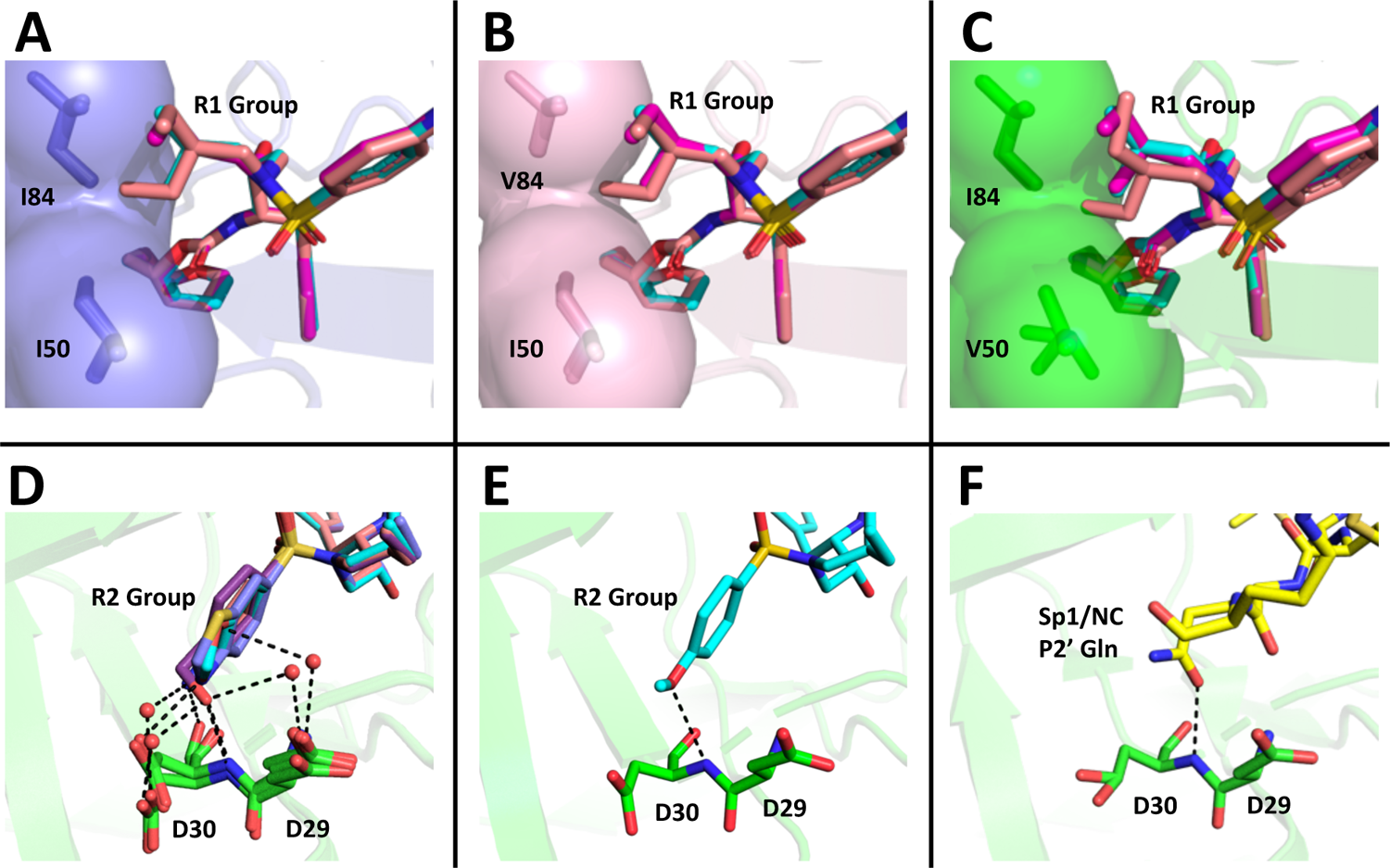
Structural Interpretation Of Protease Inhibitor Resistance and Of Residual Inhibitor Potency. **(A-C)** Hydrophobic packing in the S1’ subsite in complex with R1 structural groups; darunavir is shown in cyan, UMASS-1 in magenta, and UMASS-6 in salmon.**A)** The two forms of R1 and DRV are shown packing against wild type protease at I84 and I50 (DRV, UMASS-1, UMASS-6 in wild type protease [PDB: 6DGX (cyan), 6DGY (magenta), 6DGZ (salmon)]. **B)** Those same inhibitors packing against the I84V mutant [PDB: 6DH0 (cyan), 6DH1 (magenta), 6DH2 (salmon)]. **C)** Those same inhibitors packing against the I50V protease variant [PDB:, 6DH6 (salmon), 6DH7 (magneta), 6DH8 (salmon)]. **(D-F)** Binding interactions with Asp29 and Asp30 of the protease S2’ subsite in complex with R2 structural groups. Inhibitor/enzyme interactions are shown with black dashed lines representing hydrogen bonds (<3.0 Å). **D)** UMASS-1, UMASS-3, UMASS-4, and UMASS-5 in WT protease [PDB: 3O99 (salmon), 3O9B (purple), 309C (orange), 309D (slate blue)]. Water molecules are shown as red dots. **E**) UMASS-2 in WT protease (PDB: 3O9A). **F)** SP1/NC S2’ subsite representation of peptide substrate complexed with an inactive form of wild type protease (PDB: 1KJ7) showing an interaction between the P2’ glutamine and the protein backbone at Asp29 and Asp30.

### PR2’/R2 Modification

The inhibitors with two of the R2 modifications (R2-2 and R2-5) showed greater potency against the mildly resistant proteases (Fig. S2) and against the wild type virus (Table 1). This effect was also evident as a trend for inhibitors with the R2-2 group against the highly resistant proteases (Fig. 4E) and the highly resistant viruses (Fig. 5E). When we examined the interactions between protease and inhibitor there is a clear structural explanation (Fig. 6D-F). The less potent inhibitors interact with the protease side chains and backbone at D29 and D30 through a network of water molecules. In contrast, with the more potent R2-2 group the water molecules are replaced with direct interactions between inhibitor and protease in the S2’ subsite (Fig. 6E). This direct interaction more closely mimics the interaction made by the glutamine P2’ side chain in the optimized protease cleavage site at the SP1/NC boundary (Fig. 6F). Thus the ability to replace the water-mediated interactions with a more rigid framework that is interacting with the protease backbone deep in the S2’ pocket appears to be a unique feature of the inhibitors with R2-2. Improved potency in another series of PIs was also reported for this R2-2 P2’ moiety (Zhu et al., 2020).

### Protease Mutation Networks In the Resistant Pathways

In most of the selections either the I84V or the I50V mutation was largely fixed by the time the inhibitor concentration reached 10 nM in the culture (Fig. 2, Fig. S6). The viral cultures went through cyclical changes in population diversity as subsequent mutations were added (Fig. 2A). Because of the large number of selections it was possible to pool data and see trends in the way mutations were added in these two pathways. Resistance mutations that were detected in multiple selections summed to at least 16 positions (Fig. 1A). While these mutations are well known, their relationships to each other, and as a function of selective pressure, are less well understood. The data in Fig. 3 show strong linkage between I84V and V32I, while I50V shows linkage with I47V. Other mutations appeared often in both pathways but could be found earlier and/or more fequently in one pathway: in the I84V pathway I54L and V82I were favored; in the I50V pathway L10F, I13V, L33F, M46I, L63P, and A71V were favored. While the role of some of these changes may be in restoring fitness (Wensing et al., 2010), with a bigger fitness loss due to I50V (Henderson et al., 2012), we have demonstrate in previous studies that the mutations outside the active site have play a key role in conferring high levels of resistance (Henes et al., 2019; Ragland et al., 2017; Matthew et al., 2021; Leidner et al., 2021).

In summary, using a number of lengthy selections for resistance to a series of structurally related PIs we have been able to identify the size of the hydrophobic side chain at the P1’ position of the inhibitor as the major determinant for selection of the I84V pathway with a smaller P1’ side chain, including for DRV, versus a larger side chain driving selection of the I50V pathway. Also, one of the P2’ groups tested gave a higher level of potency even in the face of high level resistance by creating a direct interaction with the protease backbone. These changes increase the genetic barrier for the evolution of resistance and emphasize the potential utility of what a fifth generation HIV-1 PI could add to regimens with reduced drug complexity.

## Materials and Methods

### Cell lines and viruses

CEMx174 cells were maintained in RPMI 1640 medium with 10% fetal calf serum and penicillin-streptomycin. TZM-bl and 293T cells were maintained in Dulbecco’s modified Eagle-H medium supplemented with 10% fetal calf serum and penicillin-streptomycin. A wild-type virus stock NL4-3 was prepared by transfection of the pNL4-3 plasmid (purified using the Qiagen Plasmid Maxikit) into HeLa cells. For the mixture of isogenic mutant viruses the following NL4-3 variants were created, each with a single mutation in the protease with this mixture forming the virus pool for the initiation of selection with mutant viruses: L10I, K20R, K20I, L24I, D30N, V32I, M36I, M46I, M46L, I47V, G48V, F53L, I54V, I62V, L63P, A71T, A71V, G73S, V77I, V82A, V82T, I84V, N88D, N88S, L90M, I93L (Henderson et al., 2012).

### Selections

An aliquot of 3 x 10^6^ CEMx174 cells was incubated at 37°C for 2 to 3 h with 250 µl of a virus stock generated from the HIV-1 infectious molecular clone pNL4-3. The culture volume was then brought to 10 ml with RPMI medium. Each flask received one of the following inhibitors at escalating concentrations: UMass1, UMass2, UMass3, UMass4, UMass5, UMass6, UMass7, UMass8, UMass9, UMass10, DRV and no drug (ND). After 48 h and every 48 h after, the cells were pelleted by centrifugation and 10ml of fresh medium and inhibitors were added. When the culture had undergone extensive cytopathic effect (CPE) indicative of viral replication, the supernatant medium and the cells were harvested separately and stored at −80°C. The virus-containing supernatant was used to start the next round of infection, and after several rounds at the initial concentration, the inhibitor concentration was increased 1.5-fold at each subsequent round of virus passage. The level of resistance (50% inhibitory concentration [EC50]) of the single inhibitor-selected virus pools was determined by a TZM infection assay in which the PI is added to productively infected cells and the titers of supernatant virus made in the presence of the inhibitor are determined.

### TZM Infection Assay

PI dilutions were prepared by taking 10 µM stocked and performing a 5-fold serial dilution using RPMI media (final drug concentration is 100 µM). One dilution of drug was added to each well of a 24 well plate and repeated so each virus would have a full set of dilutions. Viruses for the assay were made by seeding 3 x 10^6^ CEM cells in a 24 well plate and incubating with 250 µl of virus at 37°C for 2 to 3 h before bringing the culture to 10ml with RPMI media. After 48 h the medium was changed and repeated every 48 h after until the culture had undergone CPE. Infected CEM cells were collected and diluted so that 1ml of cells could be plated in each well containing a unique drug dilution. Then 24 hours later the virus supernatant was collected from each well followed by filtering through a 0.45 µM filter then placed in −80°C. Viruses were thawed and added to 96 well plates in triplicate. TZM-bl cells were collected and diluted to a concentration of 2×10^5^ cells/ml, 100 µl were added on top of the pre-plated viruses. Plates were kept in 37°C, 5% CO_2_ in an incubator for 48 hours. After 48 hours, the cells in the plates were lysed by removing the medium, washing two times with 100 µl PBS and then lysed with 1x lysis buffer (made from 5x Promega Firefly Lysis Buffer). Plates were frozen for at least 24 hours and then thawed for 2 hours before analyzing with Promega Firefly Luciferase Kit on a luminometer.

### DNA preparation and amplification of the protease-coding region

Total cellular DNA was isolated from infected cell pellets by using the QIAamp blood kit (Qiagen). The protease-coding domain of viral DNA was amplified by nested PCR. The PCR conditions are available on request. PCR products were purified by using QIAquick PCR purification kit (Qiagen) and directly sequenced or cloned into the pT7Blue vector (Novagen) and sequenced.

### Primer-ID Deep Sequencing of viral RNA

We used the PID protocol to prepare MiSeq PID libraries with multiplexed primers. Viral RNA was extracted from plasma samples using the QIAamp viral RNA mini kit (Qiagen, Hilden, Germany). Complementary DNA (cDNA) was synthesized using a cDNA primer mixture targeting protease (PR) with a block of random nucleotides in each cDNA primer serving as the PID, and SuperScript III RT (ThermoFisher). After 2 rounds of bead purification of the cDNA, we amplified the cDNA using a mixture of a forward primer that targeted the upstream coding region, followed by a second round of PCR to incorporate the Illumina adaptor sequences. Gel-purified libraries were pooled and sequenced using the MiSeq 300 base paired-end sequencing protocol (Illumina). The sequencing covered the HIV-1 PR region (HXB2 2648–2914, 3001– 3257).

We used the Illumina bcl2fastq pipeline for the initial processing and constructed template consensus sequences (TCSs) with TCS pipeline version 1.33 (https://github.com/SwanstromLab/PID). We then aligned TCSs to an HXB2 reference to remove sequences not at the targeted region or that had large deletions.

### Protease expression and purification

The highly mutated, resistant, protease variant genes were purchased on a pET11a plasmid with codon optimization for protein expression in *E. coli* (Genewiz). A Q7K mutation was included to prevent autoproteolysis (Rose et al., 1993). The expression, isolation, and purification of WT and mutant HIV-1 proteases used for enzymatic assays were carried out as previously described (Henes et al., 2019; King et al., 2002; Ozen et al., 2014). Briefly, the gene encoding the desired HIV-1 protease was subcloned into the heat-inducible pXC35 expression vector (ATCC) and transformed into *E. coli* TAP-106 cells. Cells grown in 6 L of Terrific Broth were lysed with a cell disruptor twice, and the protein was purified from inclusion bodies (Hui et al., 1993). Inclusion bodies, isolated as a pellet after centrifugation, were dissolved in 50% acetic acid followed by another round of centrifugation at 19,000 rpm for 30 minutes to remove insoluble impurities. Size exclusion chromatography was carried out on a 2.1-L Sephadex G-75 Superfine (Sigma Chemical) column equilibrated with 50% acetic acid to separate high molecular weight proteins from the desired protease. Pure fractions of HIV-1 protease were refolded using a 10-fold dilution of refolding buffer (0.05 M sodium acetate at pH 5.5, 5% ethylene glycol, 10% glycerol, and 5 µM DTT). Folded protein was concentrated to 0.5–3 mg/ml and stored. The stored protease was used in K_M_ and K_i_ assays.

### Enzymatic assays

#### K_m_ Assay

K_m_ values were determined as previously described (Henes et al., 2019; Lockbaum et al., 2019; Matayoshi et al., 1990; Windsor and Raines, 2015). Briefly, a 10-amino acid substrate containing the natural MA/CA cleavage site with an EDANS/DABCYL FRET pair was dissolved in 8% DMSO at 40 nM and 6% DMSO at 30 nM. The 30 nM substrate was 4/5 serially diluted from 30 nM to 6 nM. HIV-1 protease was diluted to 120 nM and, and 5 µl were added to the 96-well plate to obtain a final concentration of 10 nM. Fluorescence was observed using a PerkinElmer Envision plate reader with an excitation at 340 nm and emission at 492 nm, and monitored for 200 counts. A FRET inner filter effect correction was applied as previously described (Liu et al., 1999). Data corrected for the inner filter effect was analyzed with Prism7.

#### K_i_ Assay

Enzyme inhibition constants (K_i_ values) were determined as previously described (Henes et al., 2019; Lockbaum et al., 2019; Matayoshi et al., 1990; Windsor and Raines, 2015). Briefly, in a 96-well plate, inhibitors were serially diluted down from 2000-10,000 nM depending on protease resistance. All samples were incubated with 5 nM protein for 1 hour. A 10-amino acid substrate containing an optimized protease cleavage site (Windsor and Raines, 2015), purchased from Bachem, with an EDANS/DABCYL FRET pair was dissolved in 4% DMSO at 120 µM. Using a PerkinElmer Envision plate reader, 5 µl of the 120 µM substrate were added to the 96-well plate to a final concentration of 10 µM. Fluorescence was observed with an excitation at 340 nm and emission at 492 nm and monitored for 200 counts. Data was analyzed with Prism7.

Data will be submitted to the NIH short read archive.

## Acknowledgements

This work was supported by grants from NIGMS (P01-GM109767, R01-GM135919) and NIAID (R01-AI140970) of the NIH. This research received infrastructure support from the UNC CFAR (P30-AI050410), and the UNC Lineberger Comprehensive Cancer Center (P30-CA016086). We also acknowledge receipt of reagents from the NIH HIV Reagent Program: TZM-bl cells (J. Kappes and X. Wu contributors) and CEMx174 cells (P. Cresswell contributor). The support of the UNC High Throughput Sequencing Facility is also acknowledged.

## Author Contributions

ES, S-KL, SZ, GJL, MH, EAN, AS, and KK carried out the experiments. ES, GJL, AA, NKY, CAS, and RS analyzed the data. ES, NKY, CAS, and RS wrote the manuscript.

---

**Fig. S1.**
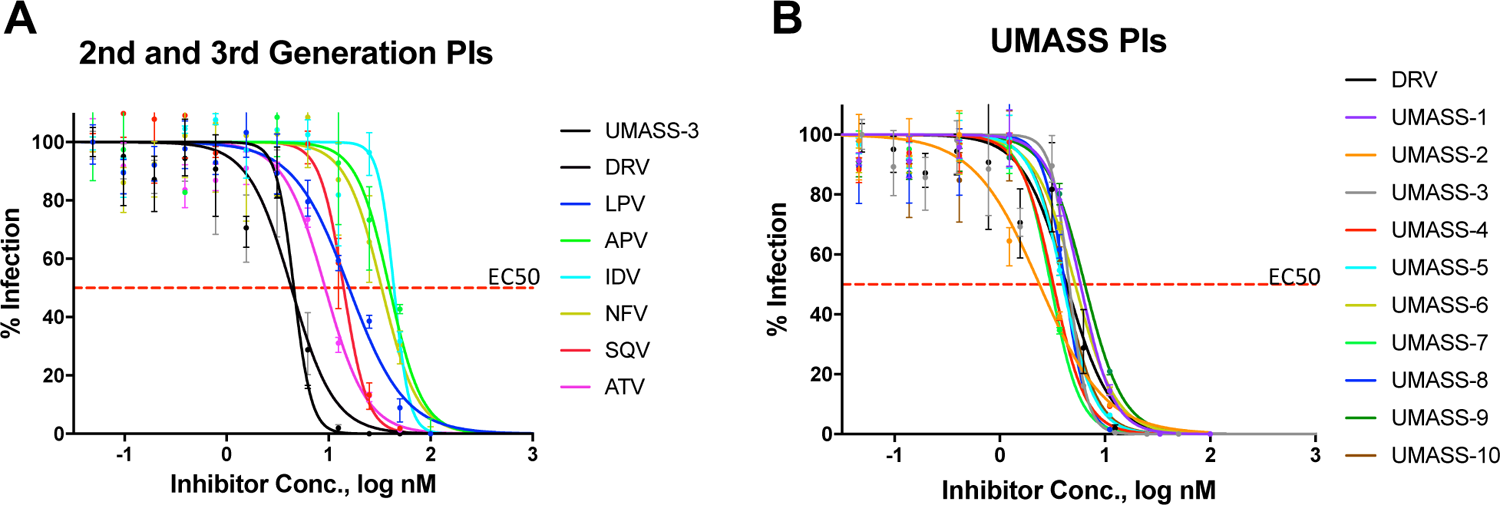
EC50 Inhibition curves for **A)** old and new protease inhibitors and **B)** fourth generation analogs of DRV (UMass 1 – UMass 10).

**Fig S2.**
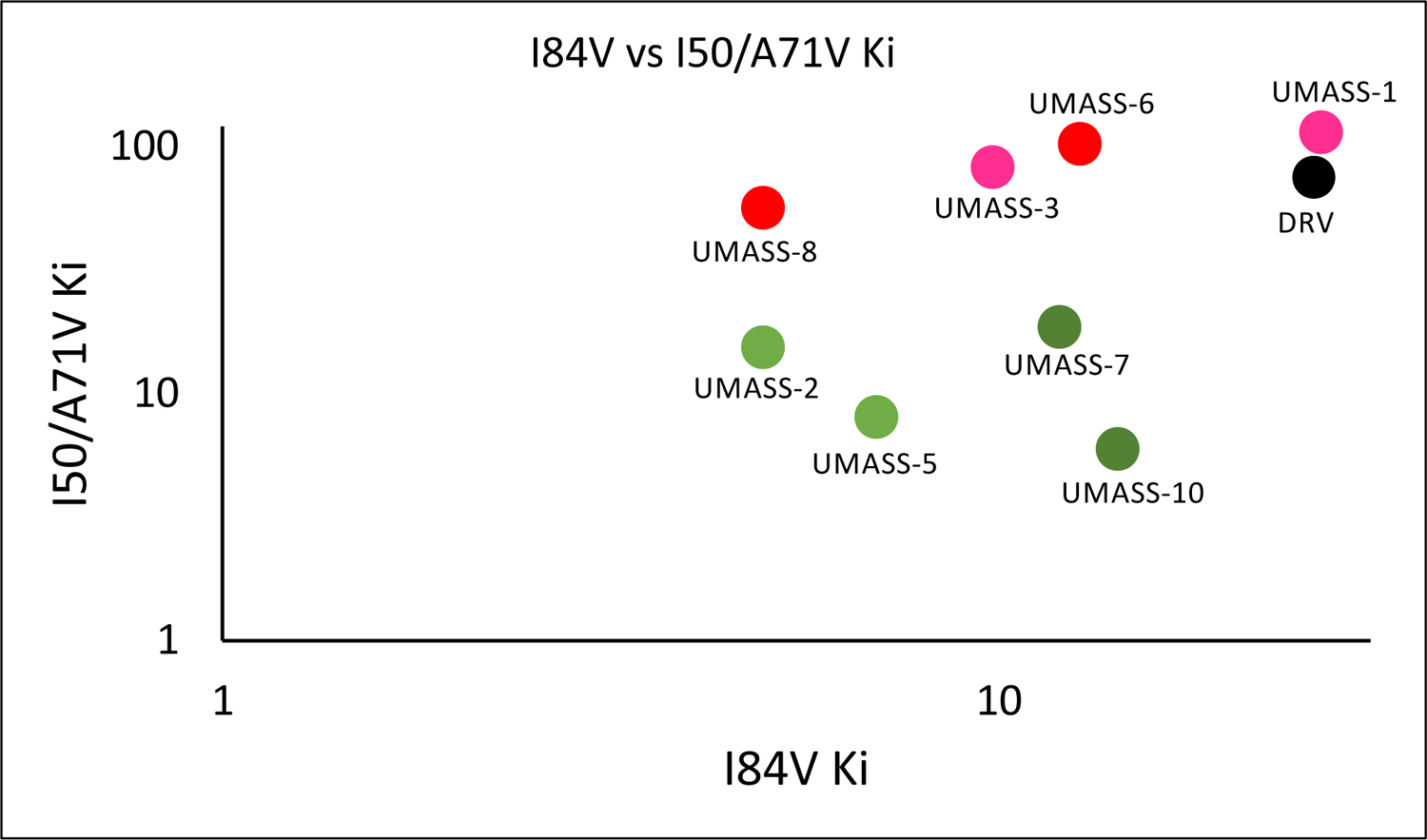
Enzymatic inhibition constant (Ki) values for UMASS-1-10 in the presence of pro with I84V or I50V/A71V mutations. UMASS-1, −3, −6, −8 show less potency towards these variants and is shaded in a red tint matching R2 groups. UMASS-2, −5, −7, −10 show increased potency towards these variants and is shaded in a green tint matching R2 groups

**Fig. S3.**
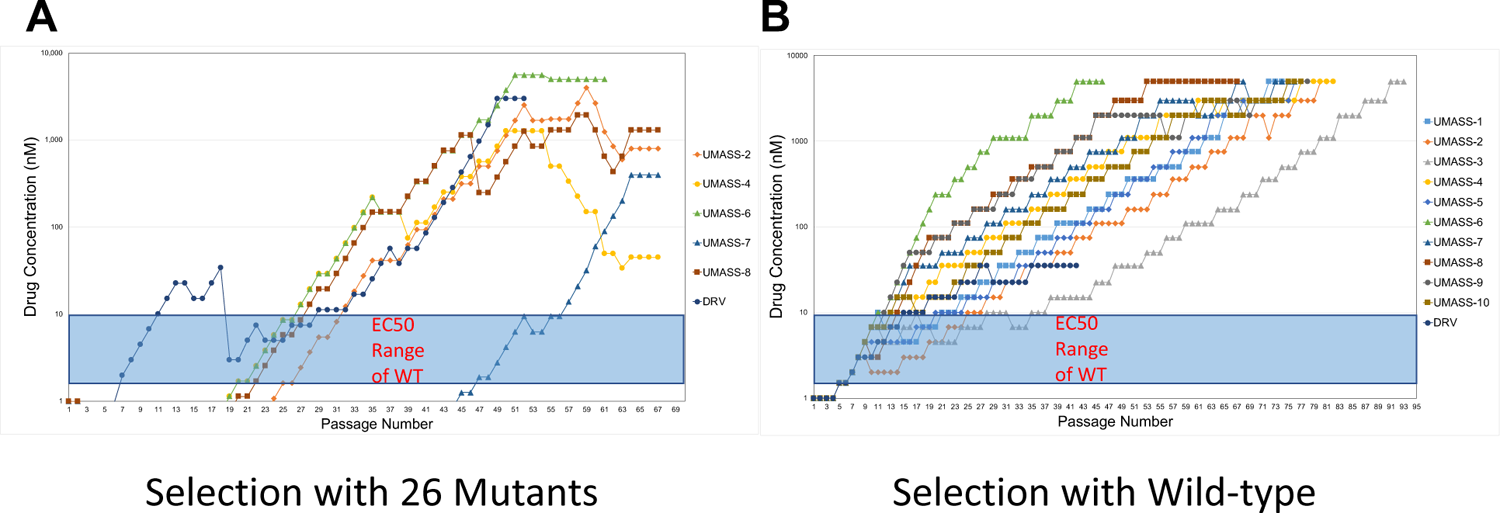
(A-B) Viral selections with DRV and UMASS-1-10 have increasing inhibitor concentrations during passaging. Passages are increased when extensive CPE is observed during PI selection. Virus generated from the **A)** 26 single mutants or **B)** NL4-3 molecular clone was passaged in the presence of increasing inhibitor concentration as described in the text. The period of time (in days) until the virus-infected culture displayed maximal CPE (massive syncytia) was 4-7 days on average.

**Fig. S4.**
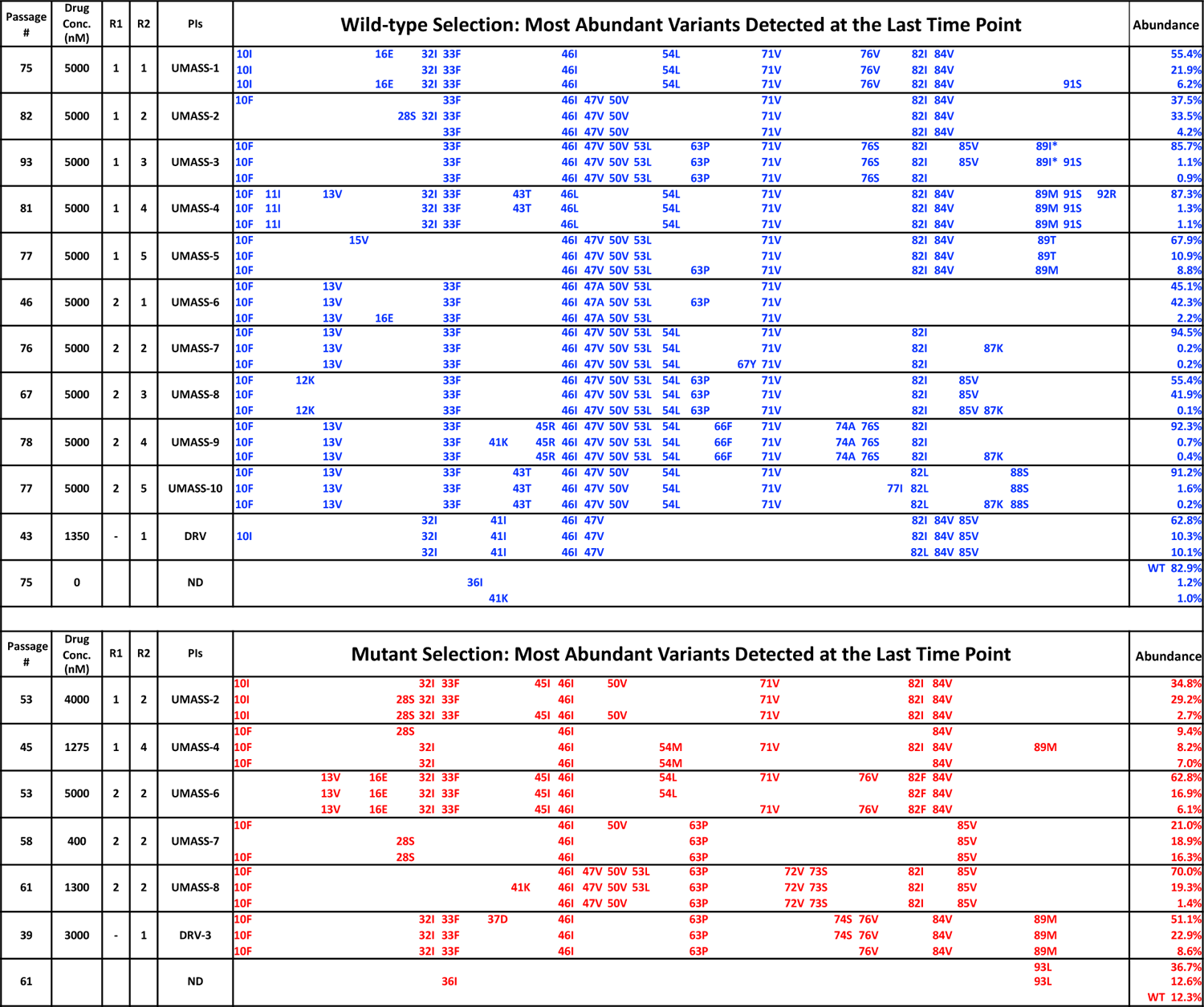
(A-B) Selection cultures for fourth generation analogs of DRV (UMASS-1 – UMASS-10), DRV and no inhibitor controls. Top 3 most abundant viral variants present at the final timepoint of each selection. **A)** Mutations arising from selections beginning with wild-type virus are shown in blue while **B)** mutations occurring in selections starting from 26-mutants are shown in red. Abundance of these variants is labeled as a percentage on the right.

**Fig. S5.**
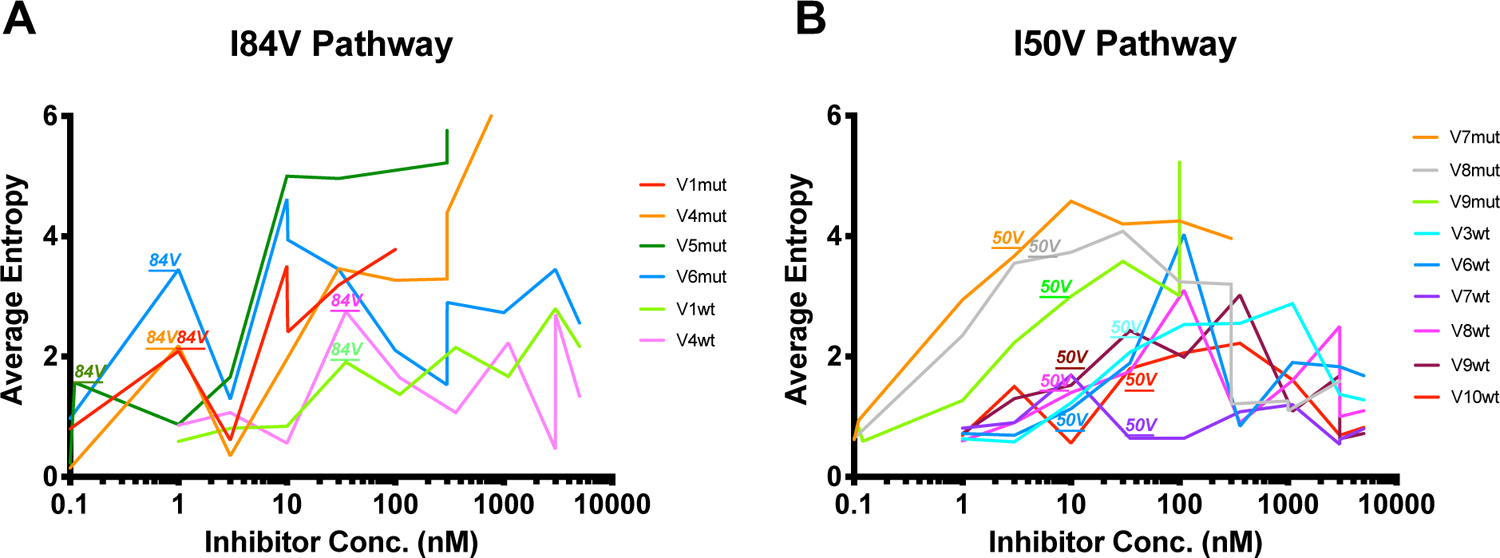
(A-B) Shannon’s entropy of viral cultures that utilize either **A)** I84V or **B)** I50V as their resistance pathway. When I84V or I50V arise in the selection cultures above the poisson distribution, that incident is labeled above the graph. I84V mutations are associated with increases/decreases in entropy while I50V mutations are not associated with major entropy differences.

**Fig. S6.**
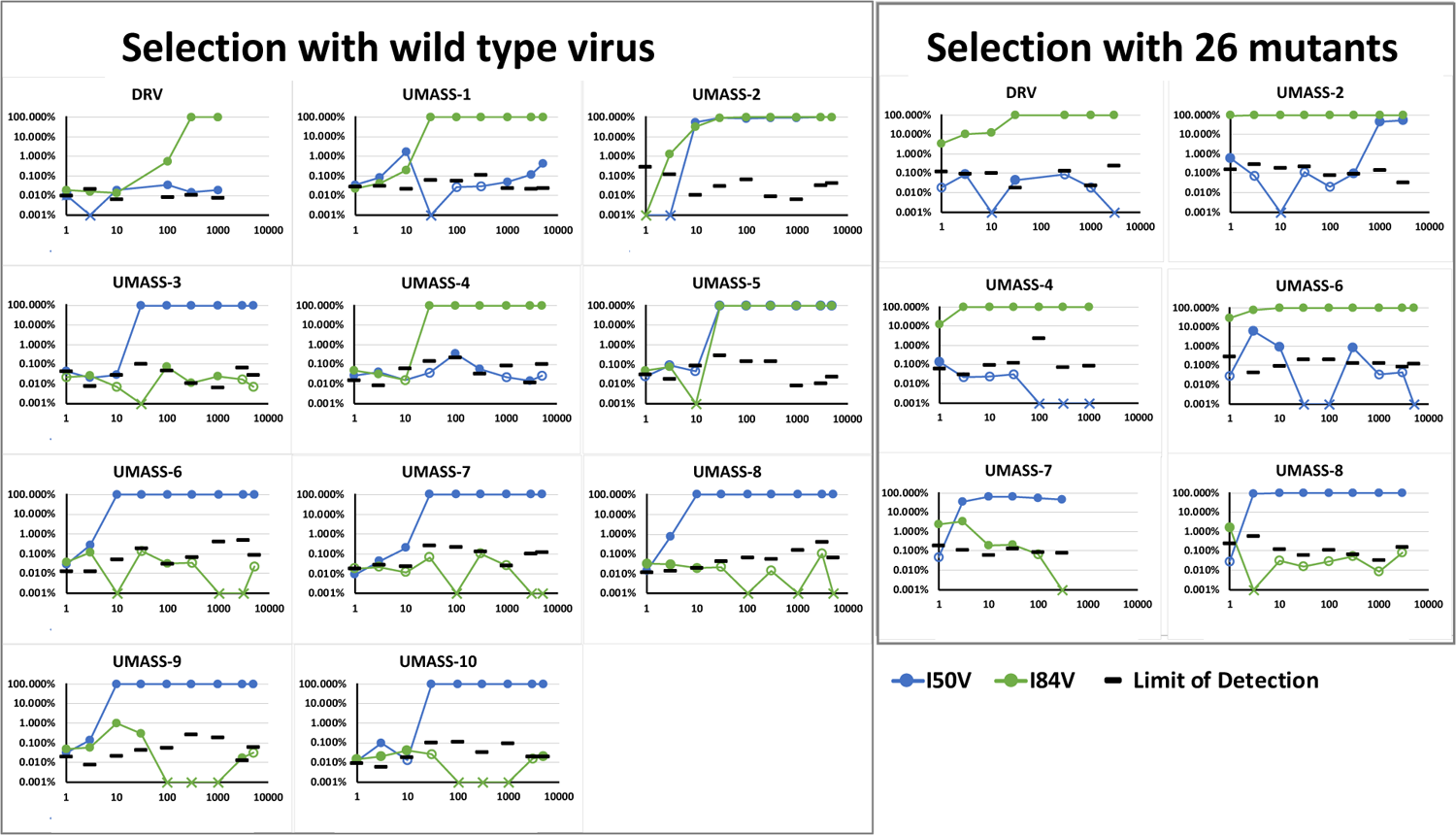
Abundance of I50V and I84V mutations at each drug concentration of all UMASS inhibitors that reached at least 1000 nM concentrations in culture. Abundance limit of detection cutoffs are shown as black lines on each timepoint to resemble our TCS poisson distribution calculation as shown in Zhou et al, 2015.

**Fig. S7.**
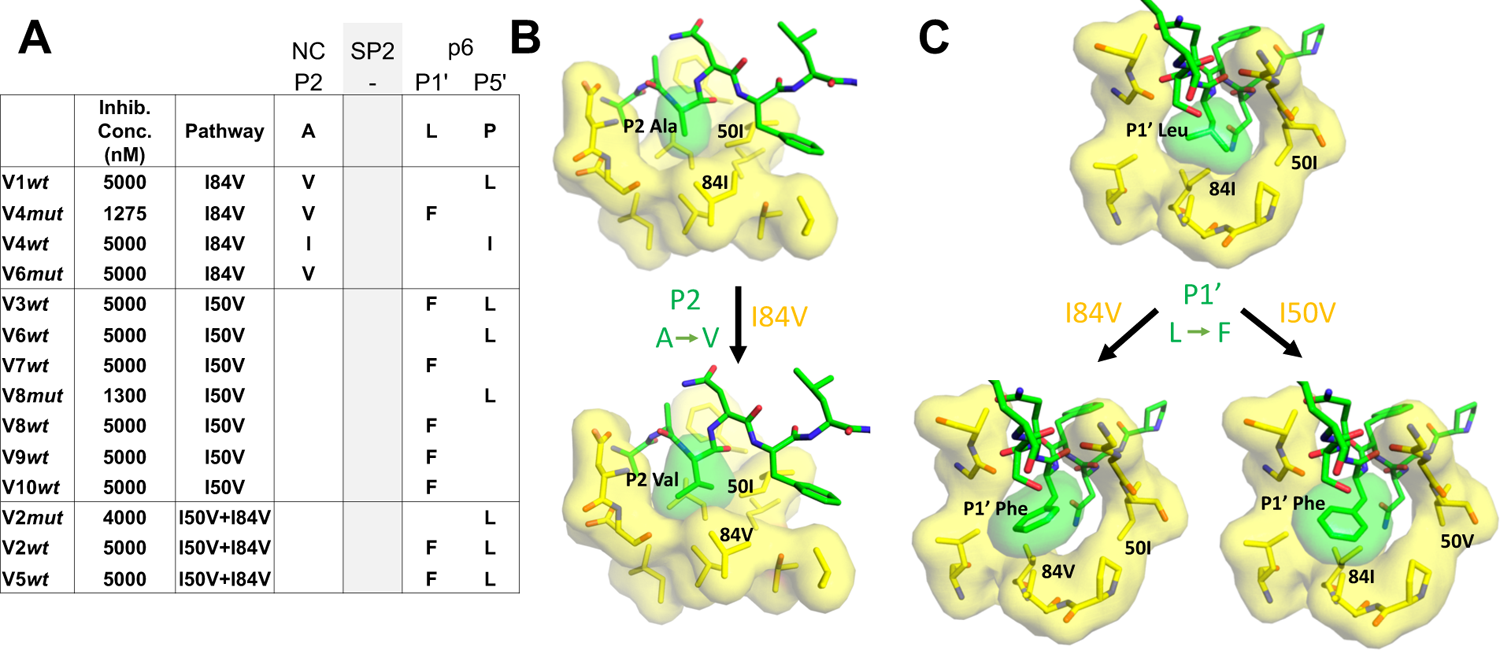
(A-B) Protease Cleavage Site Mutations Observed After Selection For High Level Resistance. **A)** Culture names, final inhibitor concentration, and resistance pathway are shown. Changes in amino acid sequence are shown for the NC/SP2 and SP2/p6 cleavage sites. **B)** P2 substitution in the NC/SP2 cleavage site with the I84V mutation modeled. Subsite structure was modeled using the original structure (PDB: 1KJH) and mutating the P2 position in the subsite pocket from Alanine to Valine. **C)** P1’ change in the SP2/p6 cleavage site with either the I84V mutation or the I50V mutations modeled. Subsite structure was modeled using the original structure (PDB: 1KJF) and mutating the P1’ position from Leucine to Phenylalanine.

