## Supplemental Figures for "Selection of HIV-1 for Resistance to Fifth Generation Protease Inhibitors Reveals Two Independent Pathways to High-Level Resistance"

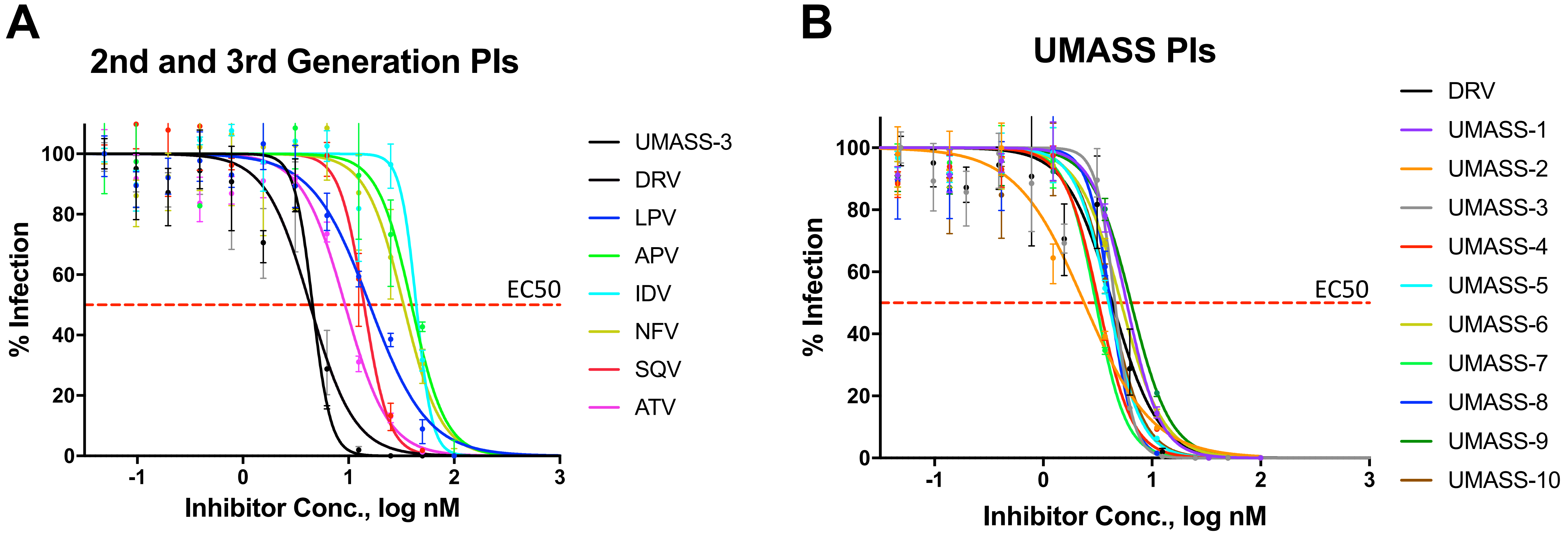


### Supplemental Figure 1

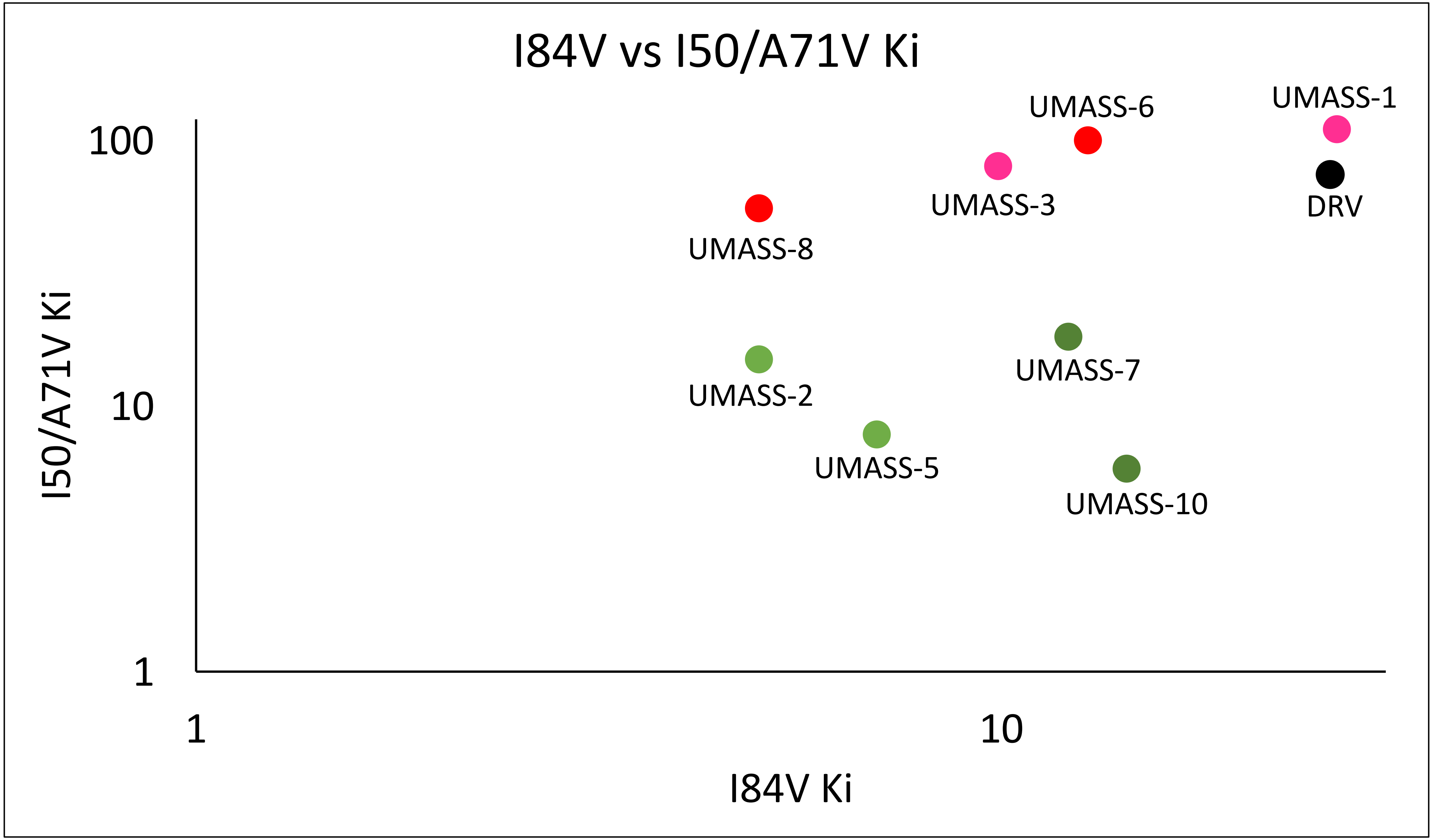


**Supplemental Figure 2**

A

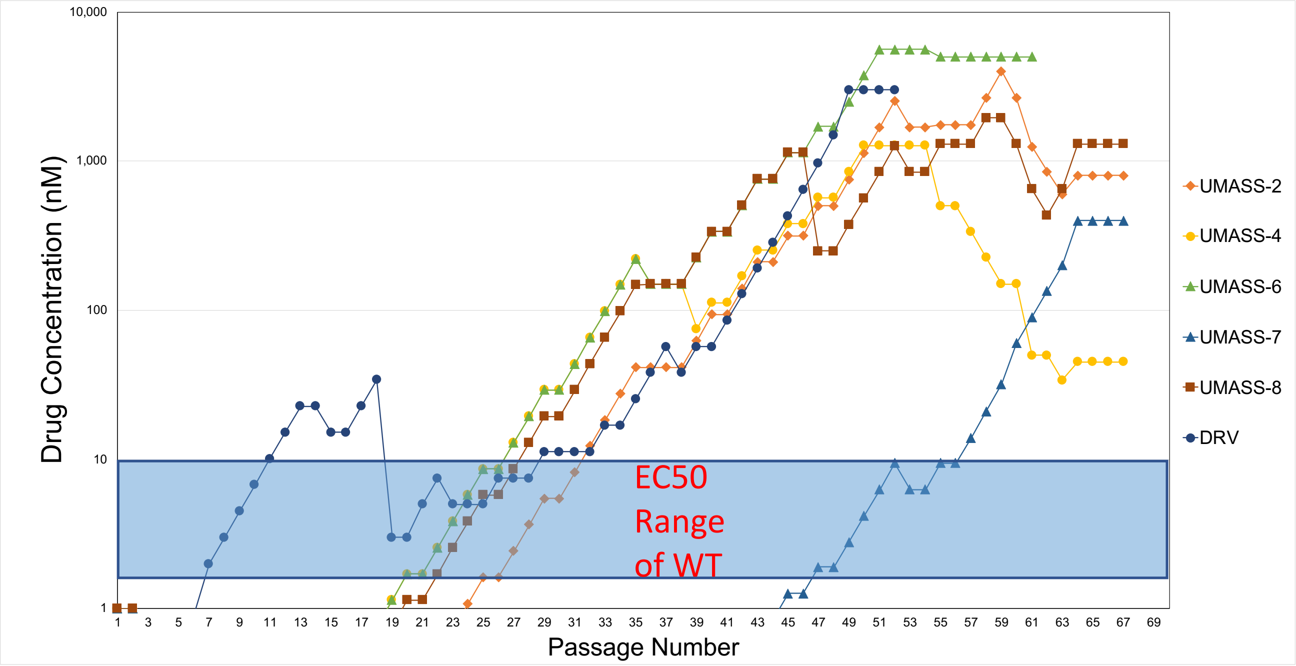


B

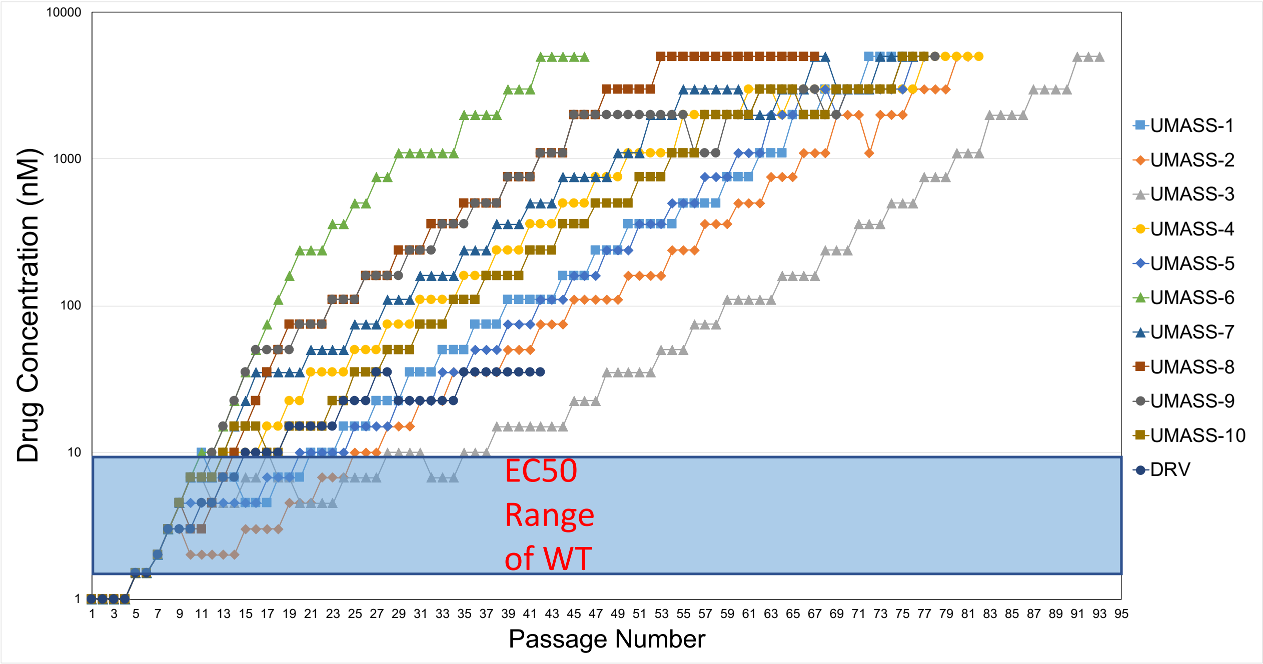


Selection with 26 Mutants

Selection with Wild-type

**Supplemental Figure 3**

| Passage # | Drug Conc. (nM) | R1 | R2 | PIs | Wild-type Selection: Most Abundant Variants Detected at the Last Time Point | Abundance |
|---|---|---|---|---|---|---|
| 75 | 5000 | 1 | 1 | UMASS-1 | 10I 16E 32I 33F 46I 54L 71V 76V 82I 84V | 55.4% |
| | | | | | 10I 32I 33F 46I 54L 71V 76V 82I 84V | 21.9% |
| | | | | | 10I 16E 32I 33F 46I 54L 71V 76V 82I 84V 91S | 6.2% |
| 82 | 5000 | 1 | 2 | UMASS-2 | 10F 33F 46I 47V 50V 71V 82I 84V | 37.5% |
| | | | | | 28S 32I 33F 46I 47V 50V 71V 82I 84V | 33.5% |
| | | | | | 33F 46I 47V 50V 71V 82I 84V | 4.2% |
| 93 | 5000 | 1 | 3 | UMASS-3 | 10F 33F 46I 47V 50V 53L 63P 71V 76S 82I 85V 89I* | 85.7% |
| | | | | | 10F 33F 46I 47V 50V 53L 63P 71V 76S 82I 85V 89I* 91S | 1.1% |
| | | | | | 10F 33F 46I 47V 50V 53L 63P 71V 76S 82I | 0.9% |
| 81 | 5000 | 1 | 4 | UMASS-4 | 10F 11I 13V 32I 33F 43T 46L 54L 71V 82I 84V 89M 91S 92R | 87.3% |
| | | | | | 10F 11I 32I 33F 43T 46L 54L 71V 82I 84V 89M 91S | 1.3% |
| | | | | | 10F 11I 32I 33F 46L 54L 71V 82I 84V 89M 91S | 1.1% |
| 77 | 5000 | 1 | 5 | UMASS-5 | 10F 15V 46I 47V 50V 53L 71V 82I 84V 89T | 67.9% |
| | | | | | 10F 46I 47V 50V 53L 71V 82I 84V 89T | 10.9% |
| | | | | | 10F 46I 47V 50V 53L 63P 71V 82I 84V 89M | 8.8% |
| 46 | 5000 | 2 | 1 | UMASS-6 | 10F 13V 33F 46I 47A 50V 53L 71V | 45.1% |
| | | | | | 10F 13V 33F 46I 47A 50V 53L 63P 71V | 42.3% |
| | | | | | 10F 13V 16E 33F 46I 47A 50V 53L 71V | 2.2% |
| 76 | 5000 | 2 | 2 | UMASS-7 | 10F 13V 33F 46I 47V 50V 53L 54L 71V 82I | 94.5% |
| | | | | | 10F 13V 33F 46I 47V 50V 53L 54L 71V 82I 87K | 0.2% |
| | | | | | 10F 13V 33F 46I 47V 50V 53L 54L 67Y 71V 82I | 0.2% |
| 67 | 5000 | 2 | 3 | UMASS-8 | 10F 12K 33F 46I 47V 50V 53L 54L 63P 71V 82I 85V | 55.4% |
| | | | | | 10F 33F 46I 47V 50V 53L 54L 63P 71V 82I 85V | 41.9% |
| | | | | | 10F 12K 33F 46I 47V 50V 53L 54L 63P 71V 82I 85V 87K | 0.1% |
| 78 | 5000 | 2 | 4 | UMASS-9 | 10F 13V 33F 45R 46I 47V 50V 53L 54L 66F 71V 74A 76S 82I | 92.3% |
| | | | | | 10F 13V 33F 41K 45R 46I 47V 50V 53L 54L 66F 71V 74A 76S 82I | 0.7% |
| | | | | | 10F 13V 33F 45R 46I 47V 50V 53L 54L 66F 71V 74A 76S 82I 87K | 0.4% |
| 77 | 5000 | 2 | 5 | UMASS-10 | 10F 13V 33F 43T 46I 47V 50V 54L 71V 82L 88S | 91.2% |
| | | | | | 10F 13V 33F 43T 46I 47V 50V 54L 71V 77I 82L 88S | 1.6% |
| | | | | | 10F 13V 33F 43T 46I 47V 50V 54L 71V 82L 87K 88S | 0.2% |
| 43 | 1350 | - | 1 | DRV | 32I 41I 46I 47V 82I 84V 85V | 62.8% |
| | | | | | 10I 32I 41I 46I 47V 82I 84V 85V | 10.3% |
| | | | | | 32I 41I 46I 47V 82L 84V 85V | 10.1% |
| 75 | 0 | | | ND | WT | 82.9% |
| | | | | | 36I | 1.2% |
| | | | | | 41K | 1.0% |

| Passage # | Drug Conc. (nM) | R1 | R2 | PIs | Mutant Selection: Most Abundant Variants Detected at the Last Time Point | Abundance |
|---|---|---|---|---|---|---|
| 53 | 4000 | 1 | 2 | UMASS-2 | 10I 32I 33F 45I 46I 50V 71V 82I 84V | 34.8% |
| | | | | | 10I 28S 32I 33F 46I 71V 82I 84V | 29.2% |
| | | | | | 10I 28S 32I 33F 45I 46I 50V 71V 82I 84V | 2.7% |
| 45 | 1275 | 1 | 4 | UMASS-4 | 10F 28S 46I 84V | 9.4% |
| | | | | | 10F 32I 46I 54M 71V 82I 84V 89M | 8.2% |
| | | | | | 10F 32I 46I 54M 84V | 7.0% |
| 53 | 5000 | 2 | 2 | UMASS-6 | 13V 16E 32I 33F 45I 46I 54L 71V 76V 82F 84V | 62.8% |
| | | | | | 13V 16E 32I 33F 45I 46I 54L 82F 84V | 16.9% |
| | | | | | 13V 16E 32I 33F 45I 46I 71V 76V 82F 84V | 6.1% |
| 58 | 400 | 2 | 2 | UMASS-7 | 10F 46I 50V 63P 85V | 21.0% |
| | | | | | 28S 46I 63P 85V | 18.9% |
| | | | | | 10F 28S 46I 63P 85V | 16.3% |
| 61 | 1300 | 2 | 2 | UMASS-8 | 10F 46I 47V 50V 53L 63P 72V 73S 82I 85V | 70.0% |
| | | | | | 10F 41K 46I 47V 50V 53L 63P 72V 73S 82I 85V | 19.3% |
| | | | | | 10F 46I 47V 50V 63P 72V 73S 82I 85V | 1.4% |
| 39 | 3000 | - | 1 | DRV-3 | 10F 32I 33F 37D 46I 63P 74S 76V 84V 89M | 51.1% |
| | | | | | 10F 32I 33F 46I 63P 74S 76V 84V 89M | 22.9% |
| | | | | | 10F 32I 33F 46I 63P 76V 84V 89M | 8.6% |
| 61 | | | | ND | 93L | 36.7% |
| | | | | | 36I 93L | 12.6% |
| | | | | | WT | 12.3% |

### Supplemental Figure 4

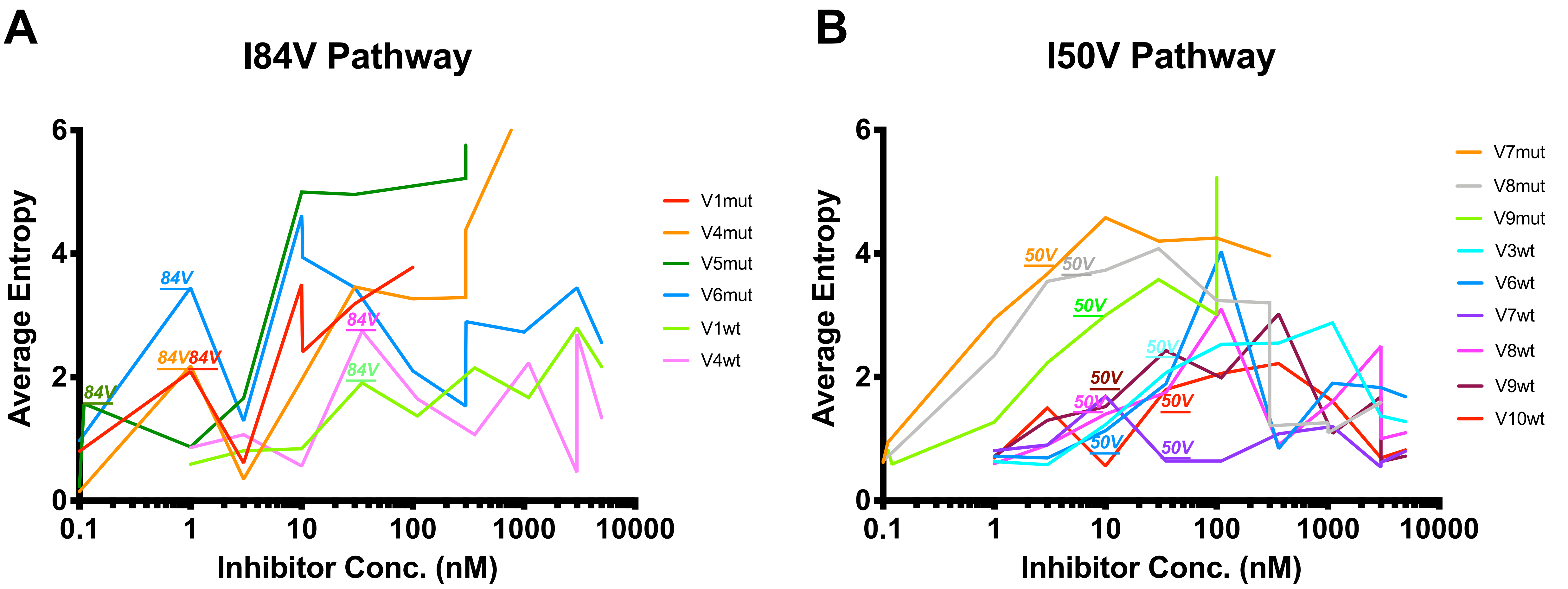


**Supplemental Figure 5**

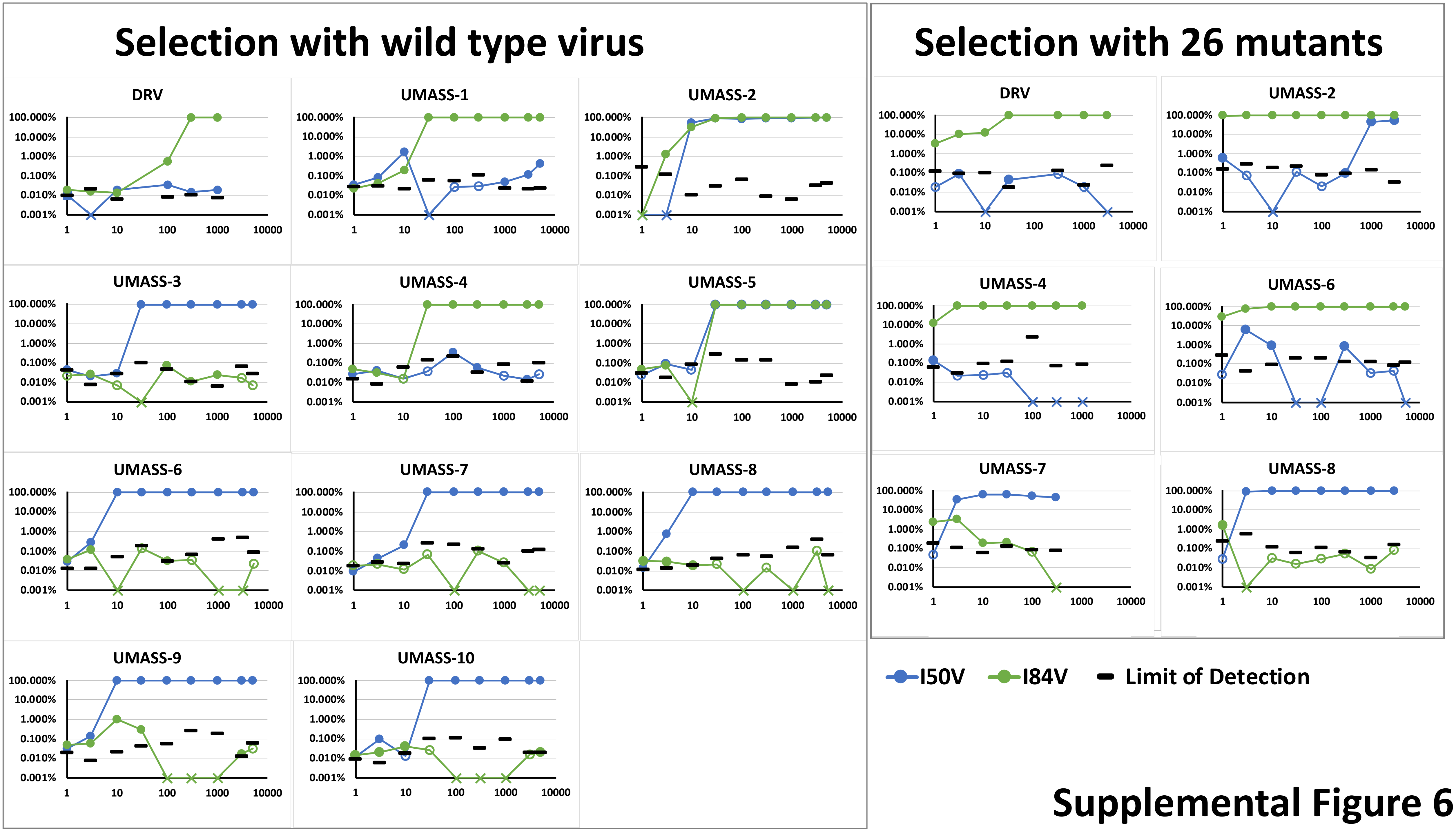


Supplemental Figure 6

| | | | NC | SP2 | p6 | |
|---|---|---|---|---|---|---|
| | | | P2 | - | P1' | P5' |
| | Inhib. Conc. (nM) | Pathway | A | | L | P |
| V1*wt* | 5000 | I84V | V | | | L |
| V4*mut* | 1275 | I84V | V | | F | |
| V4*wt* | 5000 | I84V | I | | | I |
| V6*mut* | 5000 | I84V | V | | | |
| V3*wt* | 5000 | I50V | | | F | L |
| V6*wt* | 5000 | I50V | | | | L |
| V7*wt* | 5000 | I50V | | | F | |
| V8*mut* | 1300 | I50V | | | | L |
| V8*wt* | 5000 | I50V | | | F | |
| V9*wt* | 5000 | I50V | | | F | |
| V10*wt* | 5000 | I50V | | | F | |
| V2*mut* | 4000 | I50V+I84V | | | | L |
| V2*wt* | 5000 | I50V+I84V | | | F | L |
| V5*wt* | 5000 | I50V+I84V | | | F | L |

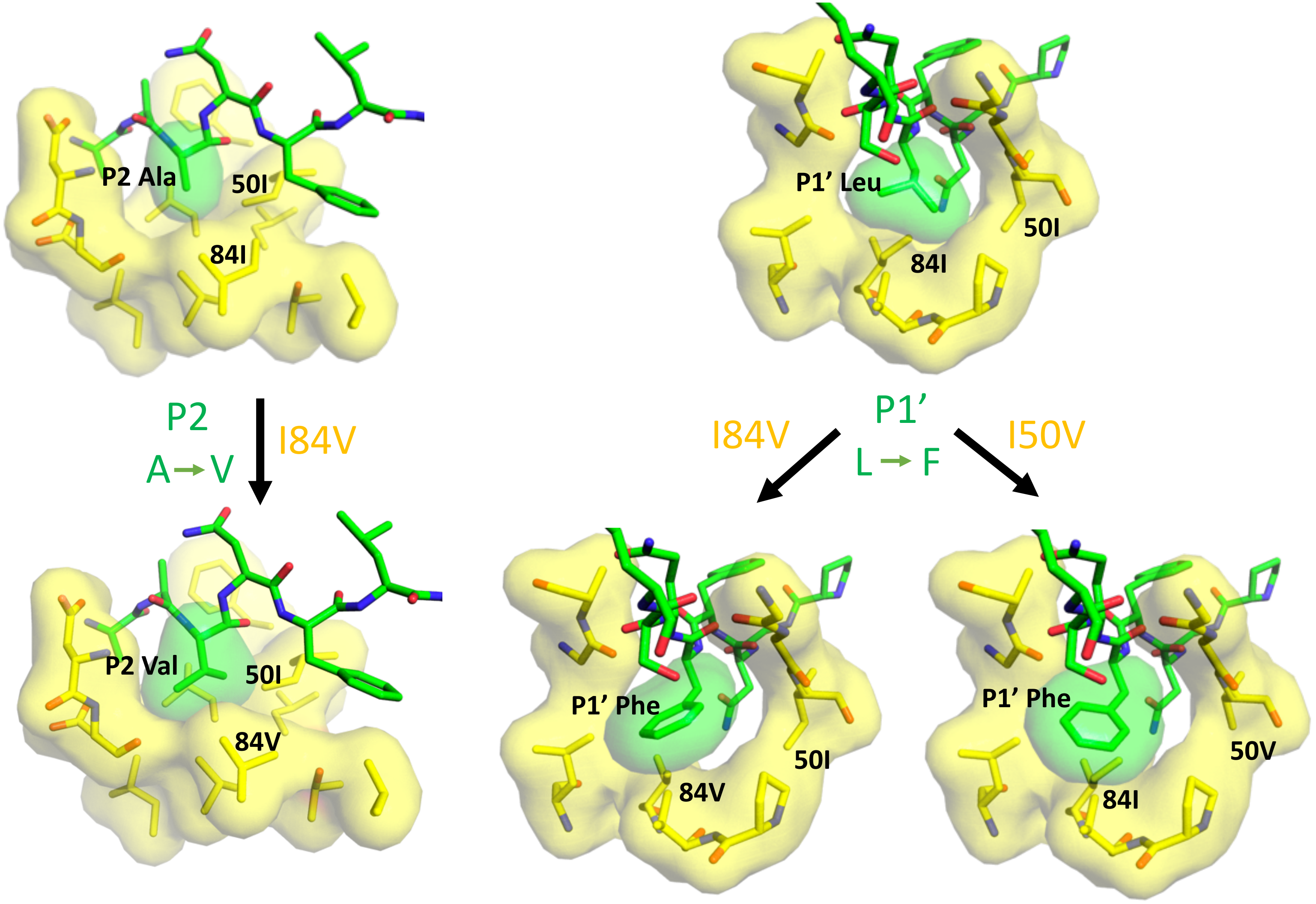


### Supplemental Figure 7
